# Exploring prokaryotic transcription, operon structures, rRNA maturation and modifications using Nanopore-based native RNA sequencing

**DOI:** 10.1101/2019.12.18.880849

**Authors:** Felix Grünberger, Robert Knüppel, Michael Jüttner, Martin Fenk, Andreas Borst, Robert Reichelt, Winfried Hausner, Jörg Soppa, Sébastien Ferreira-Cerca, Dina Grohmann

## Abstract

The prokaryotic transcriptome is shaped by transcriptional and posttranscriptional events that define the characteristics of an RNA, including transcript boundaries, the base modification status, and processing pathways to yield mature RNAs. Currently, a combination of several specialised short-read sequencing approaches and additional biochemical experiments are required to describe all transcriptomic features. In this study, we present native RNA sequencing of bacterial (*E. coli*) and archaeal (*H. volcanii, P. furiosus*) transcriptomes employing the Oxford Nanopore sequencing technology. Based on this approach, we could address multiple transcriptomic characteristics simultaneously with single-molecule resolution. Taking advantage of long RNA reads provided by the Nanopore platform, we could (re-)annotate large transcriptional units and boundaries. Our analysis of transcription termination sites suggests that diverse termination mechanisms are in place in archaea. Moreover, we shed additional light on the poorly understood rRNA processing pathway in Archaea. One of the key features of native RNA sequencing is that RNA modifications are retained. We could confirm this ability by analysing the well-known KsgA-dependent methylation sites and mapping of N^4^-acetylcytosines modifications in rRNAs. Notably, we were able to follow the relative timely order of the installation of these modifications in the rRNA processing pathway.

## Introduction

In the last decade, next-generation sequencing (NGS) technologies^1^ revolutionized the field of microbiology^2^, which is not only reflected in the exponential increase in the number of fully sequenced microbial genomes, but also in the detection of microbial diversity in many hitherto inaccessible habitats based on metagenomics. Using transcriptomics, important advances were also possible in the field of RNA biology^3,4^ that shaped our understanding of the transcriptional landscape^5,6^ and RNA-mediated regulatory processes in prokaryotes^7^. RNA sequencing (RNA-seq) technologies can be categorized according to their platform-dependent read lengths and necessity of a reverse transcription and amplification step to generate cDNA^8^. Illumina sequencing yields highly accurate yet short sequencing reads (commonly 100-300 bp). Hence, sequence information is only available in a fragmented form, making full-length transcript- or isoform-detection a challenging task^9,10^. Sequencing platforms developed by Pacific Bioscience (PacBio) and Oxford Nanopore Technologies (ONT) solved this issue. Both sequencing methods are *bona fide* single-molecule sequencing techniques that allow sequencing of long DNAs or RNAs^11,12^. However, the base detection differs significantly between the two methods. PacBio-sequencers rely on fluorescence-based single-molecule detection that identifies bases based on the unique fluorescent signal of each nucleotide during DNA synthesis by a dedicated polymerase^12^. In contrast, in an ONT sequencer, the DNA or RNA molecule is pushed through a membrane-bound biological pore with the aid of a motor protein that is attached to the pore protein called a nanopore (Fig. 1a). A change in current is caused by the translocation of the DNA or RNA strand through this nanopore, which serves as a readout signal for the sequencing process. Due to the length of the nanopore (version R9.4), a stretch of approximately five bases contributes to the current signal. Notably, only ONT offers the possibility to directly sequence native RNAs without the need for prior cDNA synthesis and PCR amplification^13^. Direct RNA sequencing based on the PacBio platform has also been realised but requires a customised sequencing workflow using a reverse transcriptase in the sequencing hotspot instead of a standard DNA polymerase^14^. Native RNA-seq holds the capacity to sequence full-length transcripts and first attempts have been made to use ONT sequencing to identify RNA base modifications (e.g. methylations^15,16^). ONT sequencing is a *bona fide* single-molecule technique and hence offers the possibility to detect molecular heterogeneity in a transcriptome^17^. Recently, the technology was exploited to sequence viral RNA genomes^18–22^ to gain insights into viral and eukaryotic transcriptomes^18,23–25^ and to detect RNA isoforms in eukaryotes^26,27^. However, prokaryotic transcriptomes have not been characterized on the genome-wide level by native RNA-seq approaches so far as prokaryotic RNAs lack a poly(A) tail, which is required to capture the RNA and feed it into the nanopore.

**Figure 1.**
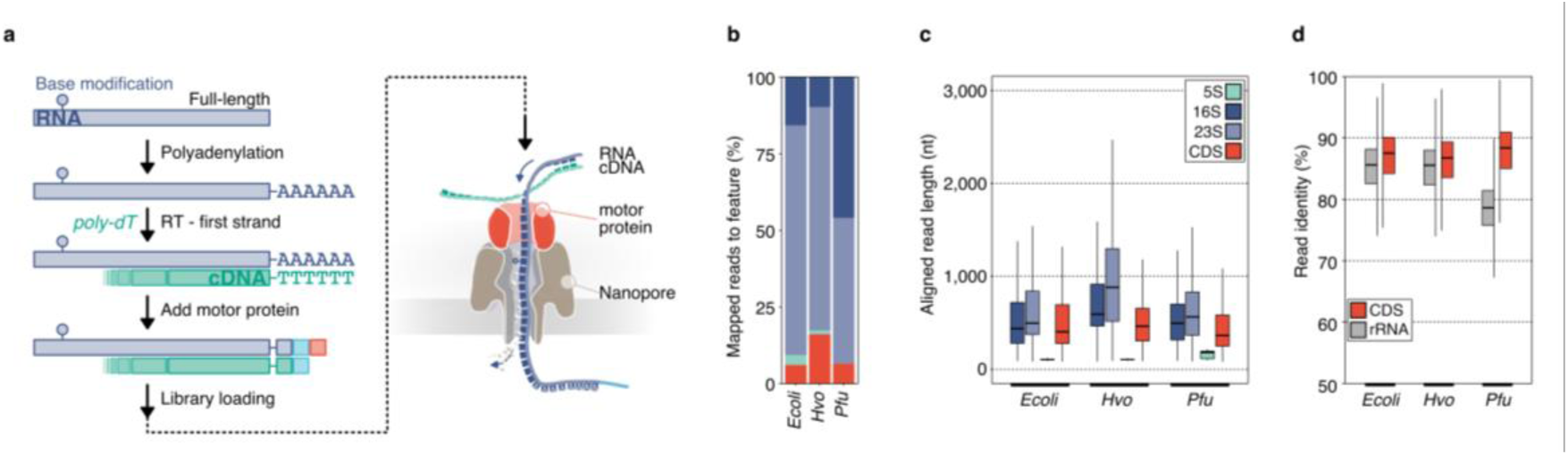
Nanopore-based native RNA sequencing of prokaryotes. **a**, Key steps of library preparation: (1) native RNA is polyadenylated, which allows library preparation using the direct RNA kit from Oxford Nanopore and sequencing on a MinION device. (2) 3’ ligation is performed to add an adapter carrying the motor-protein (red square), which unzips the RNA-cDNA hybrid and pulls the RNA through the Nanopore (detailed description see Supplementary Fig. 1a). **b**, Data sets for three prokaryotic model organisms (*Ecoli*: *Escherichia coli, Pfu*: *Pyrococcus furiosus, Hvo*: *Haloferax volcanii*) were collected and mapped to their respective reference genome. Transcript abundances of genomic features (protein coding genes (CDS): red, 5S rRNA: green, 16S rRNA: purple, 23S rRNA: light-purple) were estimated using featurecounts^50^ (TEX-treated samples are shown as example in Fig. 1). **c**, Aligned read lengths across different genomic features. **d**, Comparison of read identities between CDS (red) and rRNA (grey)-mapping reads.

Here, we present a native RNA sequencing study of bacterial and archaeal transcriptomes using Nanopore technology. We employed an experimental workflow that includes the enzymatic polyadenylation of prokaryotic transcriptomes to make them amenable for ONT’s direct RNA sequencing kit. In the first part, we evaluated the applicability of the ONT native RNA sequencing approach to survey transcriptomic features in prokaryotes and discuss weaknesses and strengths of this method. To this end, we assessed the accuracy and reliability of native RNA-seq in comparison to published Illumina-based sequencing studies of bacterial (*Escherichia coli*) and archaeal (*Haloferax volcanii, Pyrococcus furiosus*) model organisms^28–33^. The transcriptomic analysis included determination of transcript boundaries, providing, among others, insights into termination mechanisms in archaea. We moreover tested the applicability of the ONT-based native RNA sequencing approach i) to identify transcription units, (ii) to analyze pre-ribosomal RNA processing pathways and iii) to identify base modifications in (pre-)rRNAs. Despite, intrinsic limitations of the ONT-platform, we demonstrate that the long RNA reads gathered on the ONT platform allow reliable transcriptional unit assignment. Strikingly, we gained insights into the so far poorly understood ribosomal RNA (rRNA) maturation pathway in Archaea. As RNA modifications are retained when sequencing native RNAs, we explored the possibility to trace a selection of rRNA modifications in prokaryotes. Moreover, we provide data that position the relative timely order of the KsgA-dependent methylation and acetylation of rRNAs in archaea. Together, our comparative analysis suggests that rRNA modifications are more abundant in an hyperthermophilic organism.

## Material and Methods

### Strains and growth conditions

*Escherichia coli* K-12 MG1655 cells were grown in LB medium (10 g tryptone, 5 g yeast extract, 10 g NaCl per liter) to an OD_600nm_ of 0.5 and harvested by centrifugation at 3,939 x g for 10 min at 4°C.

*Pyrococcus furiosus* strain DSM 3638 cells were grown anaerobically in 40 ml SME medium^34^ supplemented with 40 mM pyruvate, 0.1 % peptone and 0.1 % yeast extract at 95°C to mid-exponential phase and further harvested by centrifugation at 3,939 x g for 45 min at 4°C.

Markerless deletion of *Haloferax volcanii* KsgA (Hvo_2746) was obtained using the pop-in/pop-out procedure^35^. Deletion candidates were verified by Southern blot and PCR analyses. Full characterization of this strain will be described elsewhere (Knüppel and Ferreira-Cerca, *in preparation*). Wildtype (H26) and Δ*ksgA* strains were grown in Hv-YPC medium at 42°C under agitation as described previously^36^.

### RNA isolation

*E. coli* total RNA was purified using the Monarch® Total RNA Miniprep Kit (New England Biolabs) according to manufacturer’s instructions including the recommended on-column DNase treatment.

*P. furiosus* total RNA was purified as described previously^33^. In short, cell pellets were lysed by the addition of 1 ml peqGOLD TriFast™ (VWR) followed by shaking for 10 min at room temperature. After adding 0.2 ml 2 M sodium acetate pH 4.0, total RNA was isolated according to the manufacturer’s instructions. Contaminating DNA was removed using the TURBO DNA-free™ Kit (Thermo Fisher Scientific).

*H. volcanii* total RNA was purified using the RNeasy kit (Qiagen) according to the manufacturer’s instructions. Alternatively, total RNA was isolated according to the method described by Chomczynski and Sacchi^37^, including a DNA-removal step with RNase-free DNase I (Thermo Fisher Scientific).

The integrity of total RNA from *E. coli* and *P. furiosus* was assessed via a Bioanalyzer (Agilent) run using the RNA 6000 Pico Kit (Agilent). To evaluate the extent of remaining buffer and DNA contaminations, the RNA preparation samples were tested by performing standard spectroscopic measurements (Nanodrop One) and using the Qubit 1X dsDNA HS assay kit (Thermo Fisher Scientific). RNA was quantified using the Qubit RNA HS assay kit.

### Primer extension analysis

5’ends determination of mature 16S and 23S rRNAs from *H. volcanii* by primer extension was performed as described previously (Knüppel et al, Method in Molecular Biology *in press*). In brief, reverse transcription was performed with the indicated fluorescently labeled primers (oHv396-DY682: 5’-CCCAATAGCAATGACCTCCG; oHv622-DY782: 5’-GCTCTCGAGCCGAGCTATCCACC) and SuperScript III reverse transcriptase using 1 µg of total RNA as template. The resulting cDNAs and reference dideoxy-chain termination sequencing ladder reactions were separated on a denaturing 14% TBE-Urea (6 M)-PAGE. Fluorescence signals (700nm and 800nm) were acquired using a Li-COR Odyssey system.

### *In vitro* transcription assays

RNA polymerase from *P. furiosus* cells and recombinant TBP and TFB were purified as described previously^38–40^. The gene encoding histone A1 (*hpyA1*) as well as the native promoter and terminator regions was used as template for transcription reactions as described in^41^.

Run-off transcription assays^42,43^ were carried out in a 25-μl reaction volume containing the following buffer: 40 mM HEPES (pH 7.5), 2.5 mM MgCl_2_, 0.125 mM EDTA, 0.25 M KCl, 20 μg/ml BSA supplied with 100 μM ATP, 100 μM GTP, 100 μM CTP, 2 μM UTP, 0.037 MBq [α-^32^P]-UTP (Hartmann Analytics) with 8.5 nM *hpy1A* template DNA, 10.5 nM RNAP, 85 nM TBP and 52 nM TFB. Reactions were incubated at 80°C or 90°C for 10 min. The radiolabeled products were extracted with phenol/chloroform and transcription products were separated on a 8%TBE-Urea (7M)-PAGE. The gel was transferred and fixed to a Whatman chromatography paper.

Gels with radioactive samples were exposed to an Imaging Plate for autoradiography. Signals derived from radiolabeled RNA transcripts were detected with FUJIFILM FLA 7000 PhosphoImager (Fuji) and analysed with Image Lab™ Software (Biorad).

### RNA treatment and poly(A)-tailing

To prevent secondary structure formation, the RNA was heat incubated at 70°C for 3 min and immediately put on ice before TEX-treatment or poly(A)–tailing of the RNA samples. Partial digestion of RNAs that are not 5’-triphosphorylated (e.g. tRNAs, rRNAs) was achieved by incubation of the RNA with the Terminator 5’-Phosphate-Dependent Exonuclease (TEX, Lucigen). For this purpose, 10 µg of RNA were incubated with 1 unit TEX, 2 µl TEX reaction buffer (Lucigen) and 0.5 µl RiboGuard RNase Inhibitor (Lucigen) in a total volume of 20 µl for 60 minutes at 30°C. The reaction was stopped and the RNA was purified using the RNeasy MinElute Cleanup Kit (Qiagen). For *P. furiosus* and *E. coli* RNA samples, control reactions lacking the exonuclease (NOTEX) were treated as described for TEX-containing samples. In the next step, a poly(A)-tail was added using the *E. coli* poly(A) polymerase (New England Biolabs) following a recently published protocol^44^. Briefly, 5 µg RNA, 20 units poly(A) polymerase, 2 µl reaction buffer and 1 mM ATP were incubated for 15 min at 37°C in a total reaction volume of 50 µl. To stop the reaction and to remove the enzyme, the poly(A)-tailed RNA was purified with the RNeasy MinElute Cleanup Kit (Qiagen).

### Direct RNA library preparation and sequencing

Libraries for Nanopore sequencing were prepared from poly(A)-tailed RNAs according to the SQK-RNA001 Kit protocol (Oxford Nanopore, Version: DRS_9026_v1_revP_15Dec2016) with minor modifications for barcoded libraries (see Supplementary Fig. 1a). In this case, Agencourt AMPure XP magnetic beads (Beckman Coulter) in combination with 1 µl of RiboGuard RNase Inhibitor (Lucigen) were used instead of the recommended Agencourt RNAclean XP beads to purify samples after enzymatic reactions. The total amount of input RNA, the barcoding strategy and the number of flowcells used can be found in Supplementary Table 1. The efficiency of poly(A)-tailing was low. However, this could be compensated with a higher amount of input RNA. We added the control RNA (RCS, yeast enolase, provided in the SQK-RNA001 kit) to detect problems that arise from library preparation or sequencing. For the barcoded libraries, the RTA adapter was replaced by custom adapters described in https://github.com/hyeshik/poreplex and reverse transcription (RT) was performed in individual tubes for each library. After RT reactions, cDNA was quantified using the Qubit DNA HS assay kit (Thermo Fisher Scientific) and equimolar amounts of DNA for the multiplexed samples were used in the next step for ligation of the RNA Adapter (RMX) in a single tube. Subsequent reactions were performed according to the protocols recommended by ONT. The libraries were sequenced on a MinION using R9.4 flow cells and subsequently, FAST5 files were generated using the recommended script in MinKNOW.

### Data analysis

#### Demultiplexing of raw reads, basecalling and quality control of raw reads

As some bioinformatic tools depend on single-read files we first converted multi-read FAST5 files from the MinKNOW output to single-read FAST5 files using the ont_fast5_api from Oxford Nanopore (https://github.com/nanoporetech/ont_fast5_api). To prevent actual good-quality reads from being discarded (this issue was reported previously^13,45^), we included both failed and passed read folders in the following steps of the analysis. Demultiplexing was done by poreplex (version 0.4, https://github.com/hyeshik/poreplex) with the arguments --trim-adapter, --symlink-fast5, --basecall and --barcoding, to trim off adapter sequences in output FASTQ files, basecall using albacore, create symbolic links to FAST5 files and sort the reads according to their barcodes. However, to ensure consistency between non-multiplexed and multiplexed samples and because of some major improvements in the current basecalling software (guppy), albacore files were not used. Instead demultiplexed FAST5 reads and raw FAST5 reads from non-multiplexed runs were locally basecalled using Guppy (Version 3.0.3) with --reverse_sequence, --hp_correct, --enable_trimming and --calib_detect turned on. After that, relevant information from the sequencing_summary.txt file in the Guppy output was extracted to analyse properties of raw reads (see Supplementary Fig. 2, see Supplementary Table 1).

#### Mapping of reads and quantification

Files were mapped to reference genomes from *Escherichia coli* K12 MG1655 (GenBank: U00096.2)^46^, *Haloferax volcanii* (NCBI Reference Sequence NC_013967)^47^ and *Pyrococcus furiosus* DSM3638^33^ using minimap2 (Release 2.17-r941, https://github.com/lh3/minimap2)^48^. Output alignments in the SAM format were generated with the recommended options for noisy Nanopore Direct RNA-seq (-ax splice, -uf, -k14) and also with (1) -p set to 0.99, to return primary and secondary mappings and (2) with --MD turned on, to include the MD tag for calculating mapping identities. Alignment files were further converted to bam files, sorted and indexed using SAMtools^49^. Strand-specific wig and bigwig files were finally created using bam2wig (Version 1.5, https://github.com/MikeAxtell/bam2wig). To evaluate the alignments, we first calculated the aligned read length by adding the number of M and I characters in the CIGAR string^13^. Based on this, the mapping identity was defined as (1-NM/aligned_reads)*100, where NM is the edit distance reported taken from minimap2. Read basecalling and mapping metrics can be found in Supplementary Table 1. Transcriptome coverage was estimated by dividing the total number of CDS-mapping reads by the sum of all CDS genomic regions.

#### Gene expression analysis

For transcript abundance estimation we applied featureCounts (Rsubread 1.32.4) allowing that a read can be assigned to more than one feature (allowMultiOverlap = TRUE) and applying the setting for long reads (isLongRead = TRUE)^50^. Calculations were performed based on the genome coordinates of genomic feature types (tRNA, rRNA, protein-coding genes). For the abundance comparison to Illumina-sequencing, we applied a regularized log transformation from the DESeq2 package that transforms counts to a log2 scale, normalizing for the library size and minimizing differences between samples with small counts^51^ (raw count data for TEX samples in Supplementary Table 2).

#### Poly(A) tail analysis

Poly(A) tail length was estimated by nanopolish following the recommended workflow (Version 0.10.2, https://nanopolish.readthedocs.io/en/latest/quickstart_polya.html)^52^.

#### Detection of transcriptional units and annotation of transcription start sites and transcription termination sites

The definition of transcriptional units (TU) and our strategy to detect and annotate them was based on a recent study that re-defined the bioinformatical search for transcriptional units (TU)^31^. The TU annotation was performed in a two-step process in the following way: First, TU clusters were defined by collapsing all reads that overlap and fulfill certain criteria that are commented extensively in the available code for this study (https://github.com/felixgrunberger/Native_RNAseq_Microbes). In short, reads were filtered out that did not align protein-coding genes (CDS) or tRNAs, had a mapping identity below 80%, were spliced, were shorter than 50% of the gene body and did not cover either the 5’ or the 3’ untranslated region. The remaining overlapping reads were collapsed in a strand-specific manner and merged.

Finally, the collapsed reads that represent the TU cluster, were split according to the coverage drop at the 3’ region of a gene. This was achieved by calculating the sequencing depth in a window of 20 nt upstream and downstream of the corresponding TTS and applying a deliberately low threshold of 1.5x (higher coverage upstream compared to downstream, see transcriptional unit table in Supplementary Table 5).

TSS were predicted by calculating the median start position of all reads that map to one gene and cover the 5’ part of a CDS. To address the 3’ coverage bias and the underrepresentation of reads that map to the 5’end and also for the 12 missing nucleotides at the TSS in general, all reads starting at least 20 nt downstream of the annotated gene start were included. To not exclude too many reads, the position of TTS were predicted similarly, by also including reads that have end positions starting from 20 nt upstream of a gene end (TSS table in Supplementary Table 3, TTS table in Supplementary Table 4).

For the analysis of prokaryotic promoter elements, the sequences 46 basepairs upstream of the corrected transcription start site were analysed to identify relevant motifs using MEME with default options except for a custom background file, calculated from intergenic sequences of the respective organism^53^.

The analysis of terminator sequences was performed comparably by extracting all TTS that are located at the end of a TU and searching for terminators in a sequence window from −45 (upstream) to +45 (downstream) from the TTS using MEME and the custom background model. Heatmap analysis of motif positioning was performed by importing MEME FASTA information into R. Metaplots of the nucleotide enrichment analysis (compare^28,54^) were calculated by comparing the genomic sequences surrounding a TTS in a window from −45 to 45 to randomly selected intergenic positions (subsampled, n = 10000). Next, the log_2_-fold enrichement was calculated and plotted as in Fig. 2e and Supplementary Fig. 7b.

**Figure 2.**
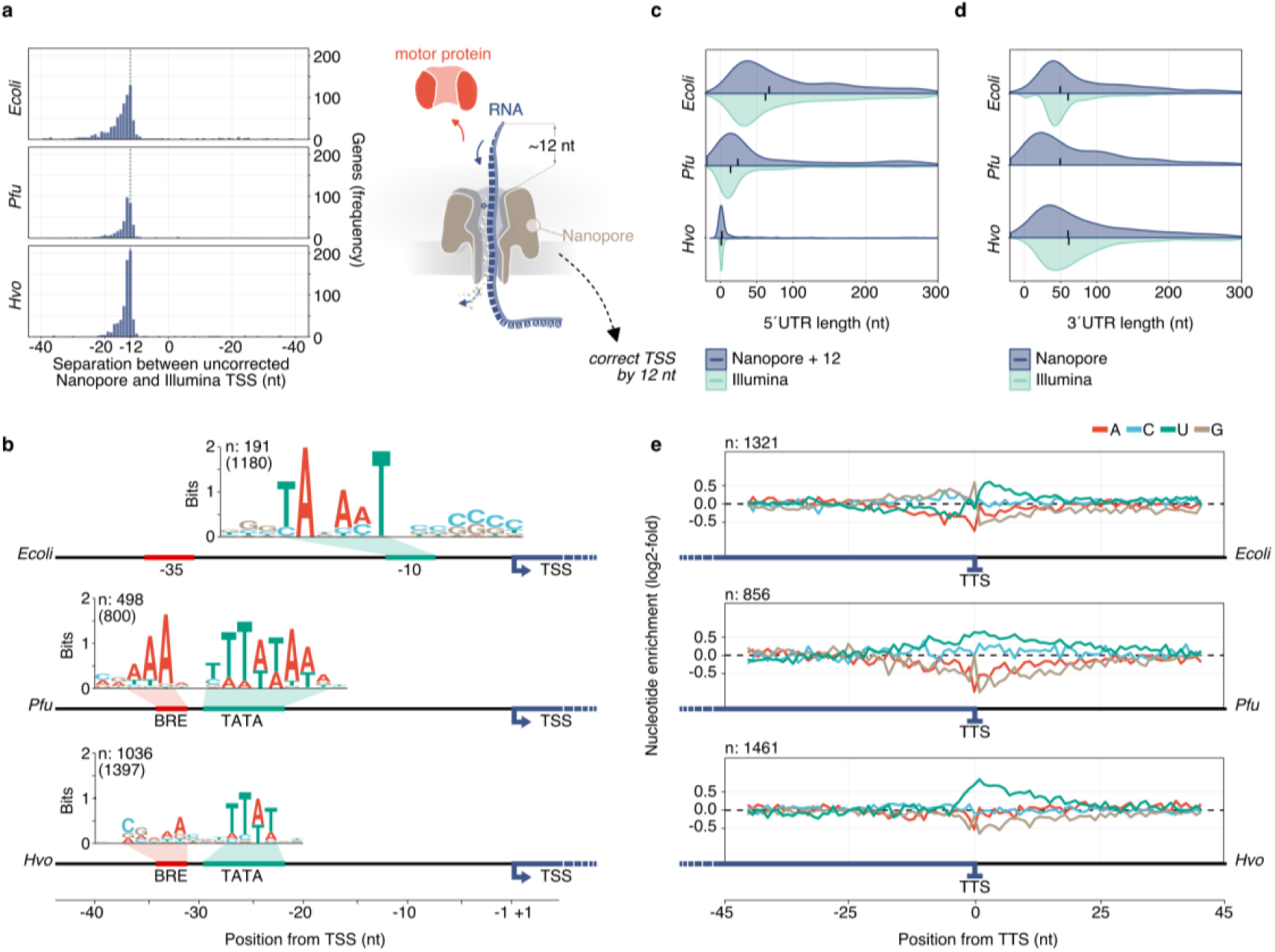
Detection of transcript boundaries. **a**, Left panel: Separation between uncorrected Nanopore-predicted TSS and comparison to Illumina d(ifferential) RNA-Seq data from published data sets for *E. coli* ^28^, *P. furiosus*^*53*^ and *H. volcanii* ^*60*^. Right panel: The translocation speed of the last 12 nucleotides (nt) is not controlled, as the motor protein is falling off. Therefore, native RNA reads are shortened by ∼12 nt. **b**, Position of TSS is corrected for 12 nucleotides to calculate the length of 5’ untranslated regions (UTR) in the Nanopore data sets (purple). 5’ UTRs are compared to d(ifferential) RNA-Seq Illumina data sets (light-green). Median values are indicated by a black bar inside the distribution (compare Supplementary Fig. 6). **c**, Length of 3’ UTRs is based on the prediction of transcription termination sites (TTS) and the comparison to annotated gene ends. Distribution of lengths is shown for Nanopore data sets (purple) and compared to Term-Seq Illumina data from *E. coli* and *H. volcanii*l (light-green)^28^. **d**, MEME analysis^53^ of extracted sequences upstream of Nanopore-predicted TSS reveals bacterial (position −10) and archaeal-specific promoter elements (BRE: B-recognition element, TATA: TATA-box recognized by transcription factor B), therefore validating the positions of predicted TSS. **e**, Nucleotide enrichment meta analysis was carried out by comparing the genomic sequences surrounding the TTS (−45 to +45) to randomly selected intergenic positions of the respective organism (n: 10000) (Terminator motifs in Supplementary Fig. 7).

RNA structural stability was predicted by folding the 45 nt long DNA upstream of the TTS using the RNAfold software from the Vienna RNA package^55^. The results were compared to randomly selected intergenic positions of the respective organism (size = 45 nt, n = 10000) and to published TTS positions derived from Term-Seq data^54,56^.

Additionaly, accuracy of TTS prediction was analysed by comparing the 3’UTRs in *H. volcanii for* genes, that were detected in both Term-Seq and Nanopore data (TEX set *H. volcanii* was used for this analysis^56^). The strength of the association between the two variables was investigated by calculating Pearson’s correlation coefficient.

#### Detection of rRNA processing sites and classification of rRNA intermediates

Processing site detection in bacteria and archaea was done by enrichment analysis of start and end positions of reads mapping to the relevant rRNA region. Next, co-occurence analysis in *E. coli* was performed by (i) categorizing reads according to enriched and literature-expected 5’ positions, (ii) selecting all reads that start within +/-1 from the relevant 5’ position and (iii) analysing the respective read ends. Note that non-circular reads were 5’extended by 12 nucleotides which corresponds to the actual transcript start. Exemplary reads of selected categories with enriched connected terminal positions were visualised in a genome browser-like view.

In addition to terminal enriched positions, read categories in archaea are based on the number of junctions that are detected (njunc argument, compare post-16S-bhb/pre-ligation and RNA chimera category in Supplementary Fig. 14), and clipping properties of the alignments on the 5’ end of the reads (see circular RNA detection).

#### Circular RNA detection and confirmation

Circular reads were initially observed in a subset of reads, which end near/at the 5’ cleavage site of the bulge-helix-bulge (bhb), but are extensively left-clipped, which happens during mapping if the nucleotides further upstream do not match the 5’ leading, but the 3’ trailing region of the rRNA. Accuracy of 5’ and 3’ cleavage site detection using Nanopore reads was further evaluated by secondary structure prediction of the potential bulge-helix-bulge regions using RNAfold^55^.

To investigate circular rRNA reads in more detail, a permuted linear sequence was created. This sequence contained 500 nt upstream of the annotated rRNA end to the predicted 3’cleavage site of the bhb site and was joined with the 5’ cleavage site of the bhb up to 500 nt downstream of the annotated rRNA start. Nanopore reads were re-mapped to the linear permuted sequence and again categorised by their 5’ ends and 3’ ends as circular (3’random breaks within the rRNA) or opened-circular (3’breaks at mature rRNA start, compare Supplementary Fig. 15). Additionally, a shorter permuted sequence was created that included x-1 nt upstream and downstream of 3’-bhb cleavage and 5’-bhb cleavage, respectively, where x is depending on the available read length of the additional Illumina data sets used (*H. volcanii*: 100 nt,; *P. furiosus*: 75 nt)^29,33^. Illumina reads were also re-mapped to the permuted sequence using bowtie2, allowing for no mismatches (-D1 -N 0 - L32 -I S,1,0.50 --score-min C,0,0) and filtering out all reads that do not overlap the joined 3’-to-5’-bulge.

#### Modified base detection

The performance of two different approaches (Tombo vs. basecalling properties) for the detection of modified bases was evaluated:

(1) We used Tombo (Version 1.5.1, https://nanoporetech.github.io/tombo) to identify modified bases based on a comparison to a theoretical distribution (*de novo* model) and based on the comparison to a reference data set (sample-compare model)^57^. Briefly, for Fig. 6f reads mapping to 16S rRNA were preprocessed, resquiggled and the raw signal plotted at a specific genomic coordinate using the respective plotting command (tombo plot genome_locations). In addition, the probability of modified bases was calculated using the detect_modification de_novo command. For Fig. 6g the signals were calculated for both samples (wildtype and deletion mutant) and compared using the *control-fast5-basedirs* and *overplot Boxplot* option. For Fig 7b reference data sets were created by sorting the reads mapping to the 16S rRNA based on the pre-determined rRNA maturation categories. 5’-extended pre-rRNA were used in all cases as a background data set in the sample-compare approach. Probabilities were calculated for the sample-compare model for all read categories and plotted using custom R-scripts.

**Figure 3.**
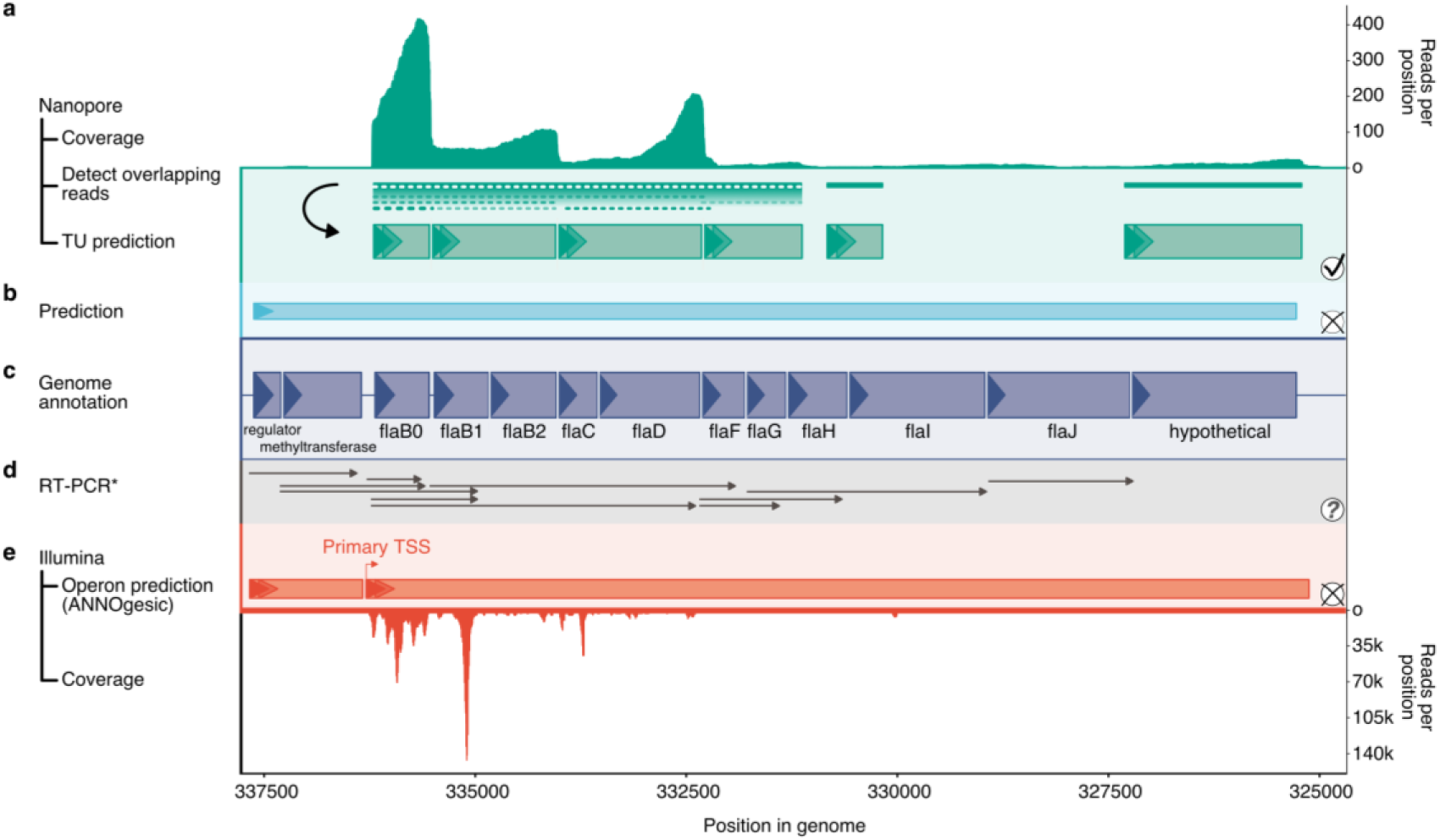
Transcription unit (TU) annotation of the flagellum-operon in *P. furiosus*. **a**, Coverage of Nanopore reads is shown in the top panel. TU prediction is performed by detection and linkage of overlapping reads and splitting them according to a 3’ drop in coverage (see Supplementary Fig. 8). Predicted TUs are drawn with green boxes according to scale. **b**, Comparison to bioinformatical prediction using the DOOR2 database^60^. **c**, Genome annotation with abbreviated gene names, boxed drawn to scale and strand indicated by triangles^33^. **d**, Comparison to results from published RT-PCR experiments^78^. All transcripts detected are drawn by arrows. **e**, Operon prediction based on mixed Illumina-Seq (coverage in lower panel) and predicted by ANNOgesic^33,142^. The primary transcription start site (TSS) of the large transcriptional unit is highlighted.

**Figure 4.**
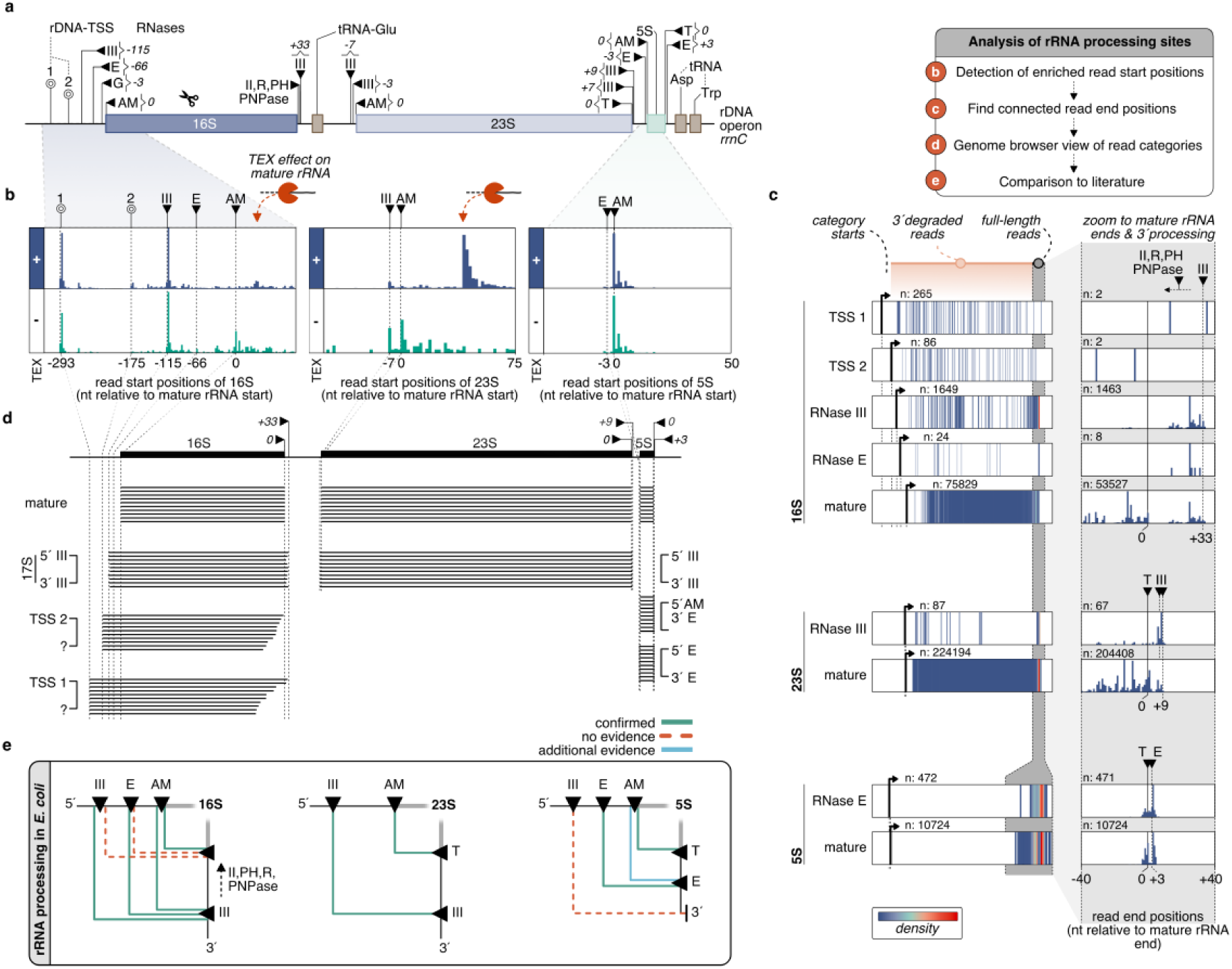
Detection and confirmation of rRNA processing sites in *E. coli*. **a**, Transcription of the rDNA locus (rrnC) is starting from two promoters (transcription start sites at −293 and −175)^84^. Precursor RNAs are cleaved by RNases (black triangles) at depicted positions^80,81,85,89^. **b**, Histograms of read start positions for 16S, 23S and 5S rRNA. Positions are relative to the annotated boundaries of mature rRNAs and shown for TEX (+, purple) and NOTEX (-, green) samples. **c**, Read start positions were used to classify categories and analyse the co-occurrence of read start to end positions. In the left panel, the color-coded density (low: blue, high: red) of read end positions from the category start position (black arrow) to the expected 3’ terminal area (grey area) is shown for the selected categories. While all lines outside of the grey area presumably represent 3’degraded reads, the full-length read end positions inside the shaded area have been analysed in more detail in the right panel (only TEX sample is shown). **d**, Based on the co-occurrence analysis of enriched read start and end positions, single reads were extracted and are visualised in a genome-browser view. **e**, The action of endo- and exonucleases (black triangles) is required for the maturation of rRNAs in *E. coli*. The multi-step maturation process leads to intermediates we could confirm (green lines) using Nanopore sequencing. While red dashed lines indicate intermediate pre-rRNA we cannot detect, blue lines indicate the presence of an additional intermediate.

**Figure 5.**
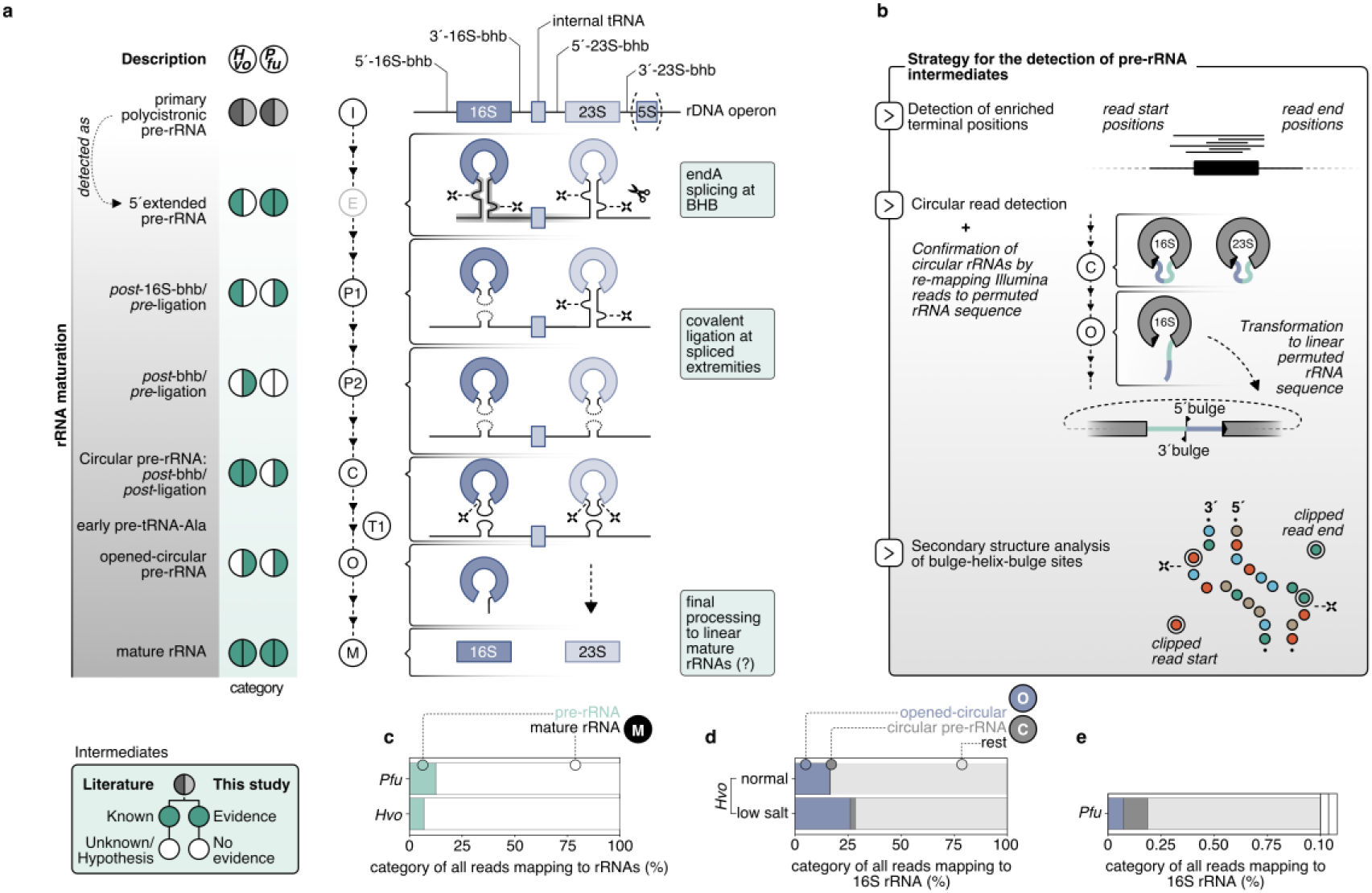
Update of the archaeal rRNA processing model. **a**, The processing stems of the primary polycistronic pre-rRNA, formed by 5’-leader and 3’-trailer sequences, contain bulge-helix-bulge motifs that are recognized and cleaved by the endonuclease endA. This is followed by the covalent ligation of the resulting extremities, which leads to archaeal-specific circular pre-16S and pre-23S rRNAs. Further maturation steps are so far unkown. The multi-step maturation process was analysed based on the strategy depicted in b and compared to already known events. **b**, Strategy for the detection of pre-rRNA intermediates: Categories were first selected based on enriched terminal positions. Clipping abnormalities lead to the detection of circular reads that could be verified by re-mapping Nanopore and Illumina reads to a linear permuted rRNA sequence containing the joined 3’-to-5’bulge region. The exact position of the joined region was additionally verified by secondary structure analysis of the bulge-helix-bulge sites^55^. **c**, Quantification of reads mapping to the mature rRNA (purple) and precursors (brown). **d**, Quantification of full-circular (green) and open-circular (orange) reads in *H. volcanii* wt and low-salt sample and in *P. furiosus*. The total number of these circular reads was compared to the number of reads mapping to the 16S rRNA.

**Figure 6.**
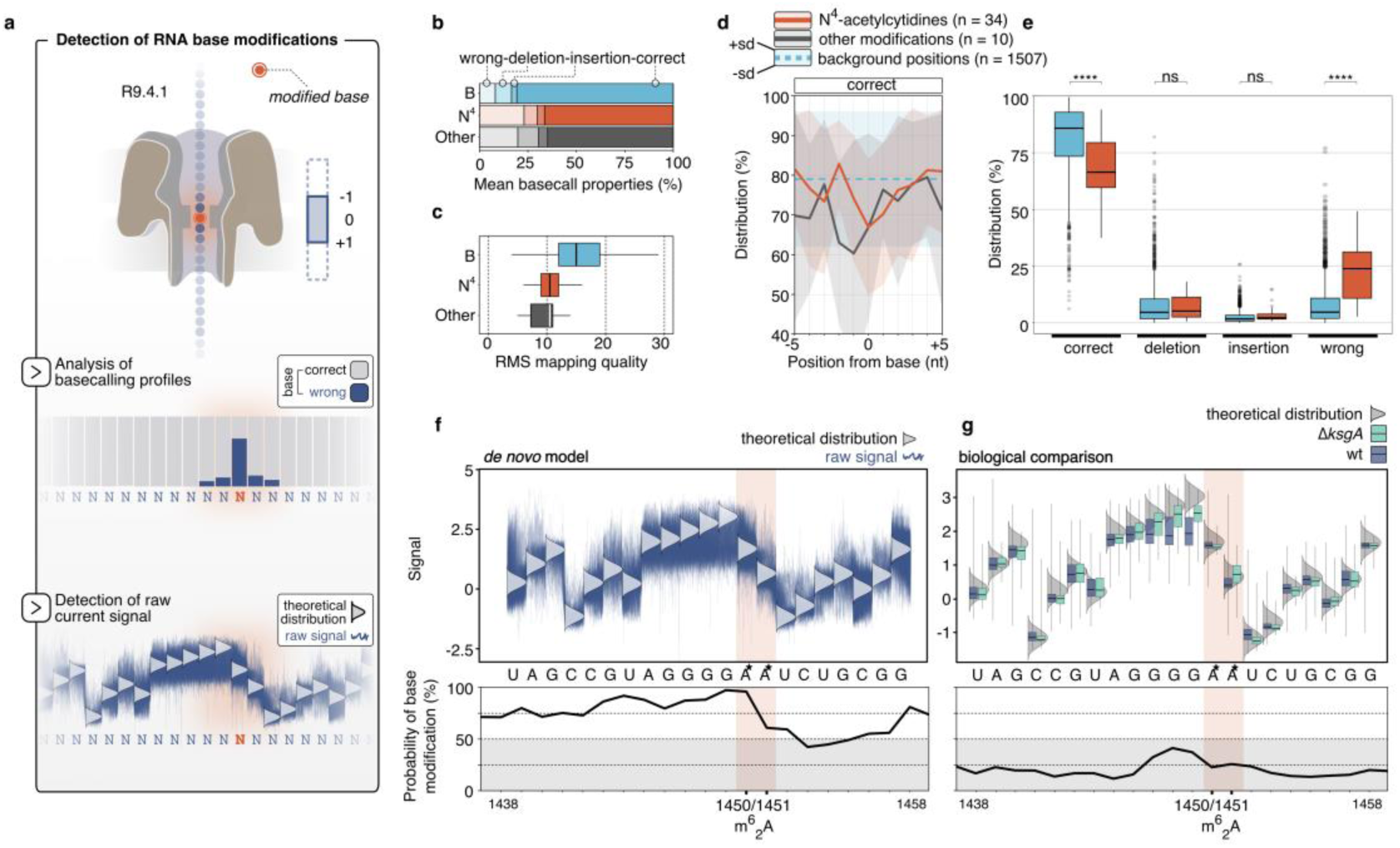
Detection of RNA base modifications in archaeal 16S rRNA based on basecalling and raw signal profiles. **a**, In the R9.4.1 pore more than one base affects the current through the pore. RNA base modifications can be predicted as the modification might alter the raw signal which can in turn influence the basecalling profile. The performance of the two strategies was evaluated for potentially present N^4^-acetylations in *P. furiosus* **(b-e**, basecalling properties) and the KsgA/Dim1-dependent dimethylation (m^6^_2_A) in *H. volcanii* **(f, g**, raw current signals). **b**, Analysis of mean basecall profiles of N^4^-acetylated positions (red), diverse other modifications (grey) and all other positions of the 16S rRNA in *P. furiosus*. The properties (wrong, deletion, insertion, correct) are shown by different transparencies. **c**, The root mean square (RMS) mapping quality gives an estimation of the overall mapping quality and is shown for the defined position categories in b. **d**, The proportion of correct basecalls is shown in a window from - 5 to +5 from the presumably modified/or background base. Shaded areas show the upper and lower standard deviation, while the lines show the mean values. **e**, Distribution of basecall properties for the 34 N^4^-acetylcytidines. Statistical significance (p-values, T-test) is indicated by asterisks (p-value > 0.05: ns (not significant), p <= 0.0001: ****). **f**, Raw signal of reads (blue squiggles) mapping to 16S rRNA in *H. volcanii* are compared to the theoretical distribution of native non-modified RNA (grey distribution) using the *de novo* detection model in tombo in the upper track.^57^ The m^6^_2_A modification at position 1450/1451 (from 16S start) is indicated by an asterisk in the sequence track. The probability of each base to be modified (in %) is calculated and shown in the lower panel for the selected sequence. **g**, Position-specific boxplot comparison of signals from sequences surrounding the m^6^_2_A modification in *H. volcanii* wildtype (blue) and the Δ*ksgA* mutant. The theoretical distribution of read signal is indicated by a grey distribution curve for every base. The probability is computed based on the comparison of the two samples.

**Figure 7.**
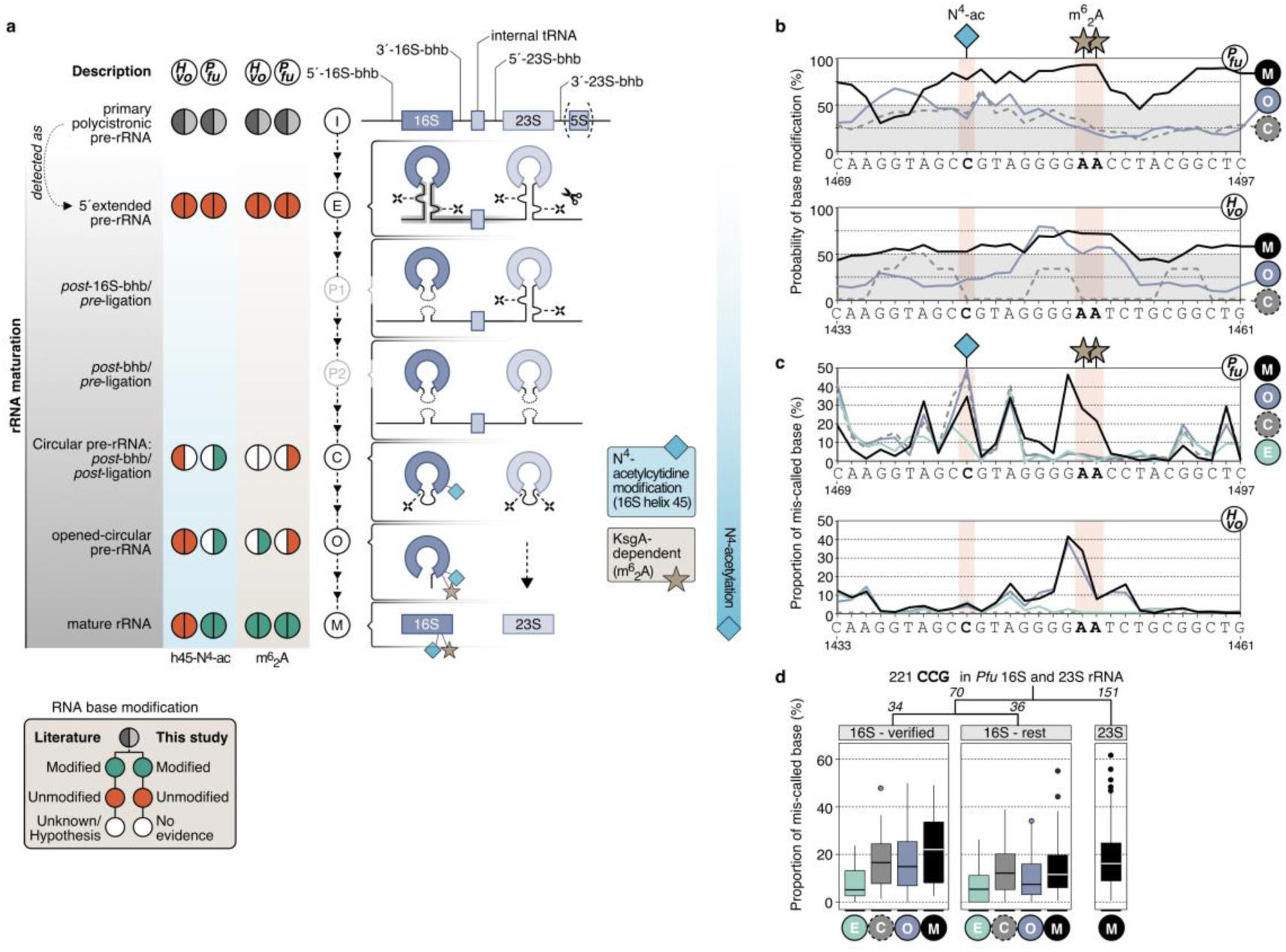
Detection of RNA base modifications at different stages of rRNA maturation in archaea. **a**, During the maturation of ribosomal rRNAs base modifications are introduced at different time points. While the the KsgA/Dim1-dependent dimethlyation (m^6^_2_A) is proposed to function as a quality control during late biogenesis (mature and open-circular pre-rRNA), N^4^-acetylations in *P. furiosus* seem to be added successively during rRNA maturation. **b**, Probability of base modifications calculated by the tombo sample-compare approach using 5’-extended pre-rRNAs as a background model for *P. furiosus* (upper panel) and *H. volcanii* (lower panel). The approach was applied to mature (M, black), open-circular (O, purple) and circular pre-rRNAs (C, grey). The illustration shows the sequence region of the 16S rRNA containing the N^4^-acetylcytidine modification in helix 45 and the dimethylation. **c**, The basecalling profile of the same section was analyed during the maturation stages. The proportion of the respective base to be mis-called (category wrong) is shown for categories M, O, C and 5’-extended pre-rRNAs (E, green). **d**, Comparison of the proportion of mis-called based for all CCGs detected in the 16S and 23S rRNA in *P. furiosus*. 34 positions have been experimentally verified in *P. abyssi* and are potentially also present in *P. furiosus*^*59*^.

(2) For calculating the frequency of correct, deleted, inserted and wrong nucleotides at a genomic position pysamstats (https://github.com/alimanfoo/pysamstats) was used. Plots were generated using custom R scripts. The results were compared to known modification sites in 16S rRNA for *H. volcanii*^58^ and *P. furiosus*. Note that the positions of modified RNA base modifications for *P. furiosus* are derived from a recently published study in *P. abyssi*^*59*^.

### Public data

In addition to the in-house generated data, we made use of other published sequencing data sets and data repositories that are described in the following.

#### Transcriptional start sites

For all three model organisms, global transcriptional start sites were mapped recently using differential RNA sequencing^29,32,33^. Position data were extracted from the Supplementary data of the publications and compared with the TSS described in the ONT data sets given that a start site was found in both data sets.

#### Transcriptional termination sites

So far there is no transcription termination data set available for *P. furiosus*. The 3’ UTR lengths of the *E. coli* and *H. volcanii* ONT sets were compared to TTS predicted based on the Term-Seq method^28,56^.

#### Transcriptional units

The widely used database DOOR2^60^ was used to compare the TU annotation for both archaeal sets. For *E. coli* a more recent, but also purely bioinformatical prediction, served as a reference set^31^.

#### Gene expression comparison

For *P. furiosus* gene abundances from ONT data were compared to fragmented RNA sequencing data of mixed growth conditions (conditions, library, sequencing, mapping described in^33^), by applying a regularized log transformation as described earlier^51^. For *H. volcanii* comparison, raw reads of a mixed RNA sequencing were extracted from the Sequence Read Archive SRA (SRR7811297)^30^ trimmed using trimmomatic^61^, (leading:20, trailing:20, slidingwindow:4:20, minlen:12), mapped to the reference genome using bowtie2 (-N 0, -L 26)^62^, converted to sorted bam files using samtools^49^ and compared to ONT data as described for *P. furiosus*. Illumina RNA sequencing data for *E. coli* were also extracted from the NCBI (SRP056485, 37°C LB), analysed as described for *H. volcanii* Illumina data and also compared to the ONT reference data.

#### Confirmation of circular rRNA precursors

For the confirmation of circular rRNA precursors we re-mapped Illumina reads to permuted rRNA sequences (see above). Illumina RNA sequencing data for *H. volcanii* (SRR3623113)^29^ and *P. furiosus* (SRR8767848)^33^ were obtained from the SRA.

## Results

### Library preparation for Nanopore native RNA sequencing of bacterial and archaeal transcriptomes

ONT allows single-molecule sequencing of RNAs in their native form. However, at present, the direct RNA sequencing kit is designed to capture polyadenylated transcripts in the first step of library preparation. As prokaryotic RNAs are not polyadenylated, we first set up a workflow that allows whole-transcriptome native RNA sequencing using the Nanopore sequencing technology (referred to as Nanopore native RNA sequencing in this work) and that can be applied to any prokaryotic organism. The key steps of the library preparation are shown in Fig. 1a: after enzymatic polyadenylation, the RNA is reverse transcribed to improve the performance by resolving secondary structures at the 3’ end (recommended by ONT)^17^. Please note that, despite the synthesis of a cDNA strand during the reverse transcription step, the RNA strand and not the DNA strand is fed into the Nanopore by the motor protein. Following this workflow, native RNAs from prokaryotic organisms can be sequenced. Depending on the necessary sequencing depth, the libraries were barcoded using poreplex (https://github.com/hyeshik/poreplex), since this is not yet supported by the official kits and protocols from Oxford Nanopore. To discriminate primary from mature rRNAs, we used a terminator exonuclease (TEX) specifically targeting 5’-monophosphorylated ends of transcripts and compared them to non-treated samples (NOTEX, see Supplementary Fig. 1a). The trimming effect of the exonuclease leads to the degradation of mature rRNAs and in turn to an enrichment of terminal positions in the non-treated samples, ultimately allowing the annotation of rRNA transcription start sites and mature rRNAs. In contrast to the experimental design of a differential RNA-seq approach, where TEX is used to detect primary transcripts in preferentially rRNA-depleted samples, we did not expect to see an effect on mRNAs, given the overall excess of rRNAs. In addition, as many Illumina sequencing-based approaches make use of a specialised library preparation design to tackle a well-defined question^8^, we evaluated the potential of native RNA sequencing to analyse multiple transcriptomic features simultaneously including the identification of *cis*-regulatory elements that govern transcription, the analysis of operon structures and transcriptional boundaries, rRNA processing and rRNA modification patterns (see Supplementary Fig. 1b).

### Sequencing yield and quality control of raw Nanopore reads

Native RNA sequencing was performed for three prokaryotic organisms: the bacterial model organism *Escherichia coli*, the halophilic archaeon *Haloferax volcanii* and the hyperthermophilic archaeon *Pyrococcus furiosus*. In order to show that native RNA sequencing can be applied to a wide variety of prokaryotic organisms, we specifically chose (i) organisms from the bacterial and archaeal domain of life with *P. furiosus* and *H. volcanii* belonging to the Euryarchaeota, (ii) organisms that are classified as mesophilic (*E. coli, H. volcanii*), hyperthermophilic (*P. furiosus*), or halophilic organism (*H. volcanii*) and (iii) organisms that differ significantly in their GC-content (*E. coli*: 50.8%^63^, *H. volcanii*: 65%^47^, *P. furiosus*: 40.8%^33^). The prepared libraries were sequenced on a MinION device and reads were collected over 48 hours on R9.4 flow cells (see Supplementary Fig. 2a). Although we did not deplete rRNAs, the total number of reads was still sufficient to also achieve good coverage of the mRNA transcriptome (*E*. coli: 9.2x, *P. furiosus*: 15.0x, *H. volcanii*: 10.3x, see Supplementary Table 1), which allowed us to perform transcriptional unit annotation and determination of transcript boundaries. Before mapping the reads to the reference genomes, the quality of the sequencing runs were evaluated based on raw read length distribution and quality of reads estimated by Guppy (see Supplementary Fig. 2b,c). To verify that no problems occurred during sequencing or library preparation the poly-adenylated spike-in control (yeast enolase) provided in the ONT-RNA kit was used. The control showed a uniform length distribution (median lengths between 1212 and 1306 nucleotides) and a very good read quality (median quality as ascertained by the Phred score between 10.8 and 12.2) in all samples, therefore, excluding any bias during sequencing (see Supplementary Fig. 2b,c, Supplementary Table 1). Lower quality in the original samples as compared to the spike-in control can be attributed to multiple reasons, including (i) compositional differences to the RNAs used to train the basecaller, and (ii) the fact that mostly ribosomal RNAs are sequenced in our samples that are known to harbor base modifications, which in turn may lead to a lower quality score especially in *P. furiosus*^64^.

### Analysis of mapped reads

An advantage of the long-read Nanopore sequencing technique is that native RNA strands can be sequenced directly as near full-length transcripts^65^. This is also reflected in the sequenced data sets as aligned lengths up to 7864 nt can be observed (Fig. 1c, Supplementary Fig. 3c). As expected, the majority of reads from all samples mapped to ribosomal RNAs, whereby the 23S rRNA represents the largest proportion (Fig. 1b, see Supplementary Fig. 3a).

In general, the read identity of CDS-mapping reads is higher than for rRNA mapping reads, but lower than the spike-in control (Fig. 1d, see Supplementary Fig. 3b,c). It is noteworthy, that accurate mapping of very short reads is currently not supported by the minimap2 mapping tool, which explains the 100 nt cut-off in our data sets (see Supplementary Fig. 3d)^17,48,66^. Unaligned reads had a median read length of 191 nt, in contrast to 572 nt for aligned reads (all data sets combined) suggesting that short reads could not be aligned properly. As small RNAs, CRISPR-RNAs or tRNAs fall below this threshold, we excluded these RNAs from further analysis. While short transcripts are problematic, longer RNAs can be sequenced and mapped accurately without loss in quality (see Supplementary Fig. 3e). As the raw read quality correlates with the mapping identity of the reads, problems during sequencing can be live-monitored in MinKNOW and the run can be canceled allowing the loading of a new library (see Supplementary Fig. 3f). Since the subsequent analysis of transcriptional units is heavily dependent on the integrity of the data, we verified the data integrity in the next steps. The addition of poly(A)_20_ (length of the reverse transcription adapter) is sufficient to allow for the annealing of the poly(T)-adapter required for reverse transcription and sequencing. This goes in line with the shortest median length we observed for the 5S rRNA (see *E. coli* TEX sample) (see Supplementary Fig. 4). For most of the transcripts, a poly(A) tail with 50 to 100 nt was detected. In addition, the overall correlation of transcript abundances calculated from sequencing data using Nanopore or Illumina technology was very high suggesting that a good coverage of the transcriptome was achieved and that native RNA sequencing is not biased towards a subset of transcripts (see Supplementary Fig. 5a,b,c, transcript abundance data in Supplementary Table 2).

### Mapping of transcriptional boundaries

#### Transcription start sites

Transcription start site (TSS) and transcription termination site (TTS) detection was based on the determination of transcriptional units (TU) (compare material and methods section)^31^. In total, we identified a comparably high number of TSS in ONT data sets compared to TSS detected by Illumina d(ifferential) RNA-seq (see Supplementary Fig. 6a)^29,32,33^. Furthermore, the substantial overlap of genes with a predicted TSS in both technolgies (see Supplementary Fig. 6a), allowed us to evaluate the accuracy of ONT TSS mapping (positions of TSS derived from ONT TEX-treated samples in Supplementary Table 3). For example, in case of *E. coli*, we could annotate the TSS for 1925 genes using the results of a published dRNA-seq study, and 1,272 TSS were detected by ONT native RNA sequencing. The portion of TSS identified only based on the ONT sequencing data (653 TSS) or Illumina sequencing data (1,436 TSS) is mostly caused by the different algorithms used and the limited sequencing depth in the ONT data sets. Strikingly, despite missing specific enrichment of primary transcripts, the median 5’ untranslated region (UTR) lengths were very similar when data from ONT native and Illumina-based RNA sequencing were compared (*E. coli*: 68 ONT vs. 62 Illumina; *P. furiosus*: 23 ONT vs. 13 Illumina; *H. volcanii*: 1 ONT vs. 0 Illumina, Fig. 2b). Please note that TSS-mapping based on Nanopore native RNA-seq data must be corrected by 12 nucleotides (Fig. 2a,b). It has been observed previously that about 12 nt are missing at the 5’ end of the sequenced RNAs. This observation can be explained by a lack of control of the RNA translocation speed after the motor protein falls off the 5’ end of the RNA (Fig. 2a)^66,67^.

Promoter analysis confirmed the presence of well-known sequence motifs of bacterial and archaeal promoters^29,33,68^. This includes the TATA-box and TFB-recognition element (BRE) characteristic for archaeal promoters and the −10 element in bacterial promoters (Fig. 2d). The - 35 element in *E. coli* has been previously shown to be less enriched compared to the −10 site^5^, which might explain why this element cannot be detected in the Nanopore data set. To analyse TSSs in more detail, we compared the 5’ UTR lengths for all genes with predicted TSS in ONT and Illumina data sets (see Supplementary Figure 6b,c,d). The overall correlation between the two techniques was high even though in some instances only a moderate correlation was found (see Supplementary Fig. 6). As expected, the correlation improves with increasing sequencing depth for a gene (>5 reads). While TEX-treatment is a common way of predicting TSS in Illumina sequencing, we observed that it is not necessary for ONT data as very similar TSS are found in both TEX and NOTEX data sets (ρ = 0.86) (see Supplementary Figure 6e).

#### Transcription termination sites

In prokaryotes, transcription termination is mediated either by a proteineous factor (Rho in bacteria^69^, CPSF in archaea^70^) or intrinsic RNA sequences (bacteria: a GC-rich sequence that forms a stem-loop followed by a U-rich sequence^71^, in archaea: poly(U) stretch^54^). Native RNA reads are sequenced in the 3’ to 5’ direction, which is a major advantage in the detection of termination sites as any bias introduced after polyadenylation can be excluded. This approach opened up the opportunity to not only map termination sites but to also gain insights into 3’ UTR lengths, for which no reference data sets for *P. furiosus* were available. The distribution of 3’ UTRs in *E. coli* and *H. volcanii* ONT data closely resembles the data from previous Illumina-based studies^28,56^. Strikingly, the length of untranslated regions at the 3’end of annotated transcripts is very similar between the three prokaryotes (Fig. 2c). In total, 1321 TTS in *E. coli*, 856 in *P. furiosus* and 1461 in *H. volcanii* were analysed (positions of TTS in Supplementary Table 4). A meta-analysis of all TTS surrounding regions revealed different sequence-dependent termination mechanisms that were confirmed using motif scanning and ΔG analysis (Fig. 2e, see Supplementary Fig. 7a-e). Our data suggest that transcription in *P. furiosus* is terminated by a double-stretch of Uridines that are distributed over a length of 22 nt, a finding that is in line with the terminator sequences detected by Term-Seq in *S. acidocaldarius*^54^ and similar to the U_(8)_ sequence in *Thermococcus kodakarensis* determined by an *in vivo* reporter assay^72^. The termination motif found in *H. volcanii* is a (U)_4_-sequence and located right after the TTS (see Supplementary Fig. 7d). In *P. furiosus*, the poly(U) is not preceded by a stem-loop structure, confirming that stem-loop structures do not play a role in hyperthermophilic organisms for general termination (see Supplementary Fig. 7e)^54,73^. However, this is less clear in *H*. volcanii, where stem loops have been shown to terminate transcripts, although less efficiently (see Supplementary Fig. 7e)^73^. The motif locations for both *Haloferax* and *Pyrococcus* ONT sets suggest that accurate TTS detection of transcripts terminated by poly(U) stretches is currently not possible. We observed that homopolymer sequences currently cannot be basecalled accurately, which leads to problems during mapping and ultimately to TTS positions that are positioned upstream of the poly(U) signal. This is also supported by a position-specific comparison of TTS in *H. volcanii* identified with Illumina and ONT reads (see Supplementary Fig. 7f,g). However, it was encouraging to see, that with increasing sequencing depth the correlation significantly improves (see Supplementary Fig. 7h). Analysing individual transcripts in *H. volcanii* and *P. furiosus*, we found that a single transcript can exhibit diverse 3’ ends. This is true for the Pilin transcript in *H. volcanii* and the Histone A1 transcript in *P. furiosus*, respectively (see Supplementary Fig. 8). Both genes are highly expressed and some transcripts carry extended 3’ UTRs. While the majority of transcripts are terminating at the first poly(U) stretch, a subset of transcripts is substantially longer and terminate at subsequent poly(U) termination signals (see Supplementary Figure 8). Interestingly, homogeneous short poly(U) signals are found both at the canonical termination site and the termination site of the elongated 3’UTR in the case of the Pilin transcript in *H. volcanii*. The same applies to the termination of the histone and Alba gene transcripts in *P. furiosus* (see Supplementary Fig. 8). The histone transcript has already been shown to terminate at four consecutive U-stretches (U1-U4) consisting of at least five U’s *in vitro*^41^. While we could confirm that the archaeal RNAP mainly terminates at the U1 site, the downstream sites seem to deviate from the three U_5_ backup TTS *in vivo*. Instead, termination already occurred at two U_4_ stretches, each upstream of U2 and U3, respectively (see Supplementary Figure 8d-f). Surveying the heterogeneity of the transcripts with an extended 3’UTR, we found a heterogeneous distribution in the length of the transcripts for both, the Histone and Alba mRNA. This pattern suggests that either a step-wise trimming of the 3’UTR occurs that eventually yields the mature RNAs or that the RNA polymerase reads through the first termination sequence, stochastically stops transcription after the first termination sequence and RNA polymerases that continued transcription beyond the first TTS terminate at one of the following TTS.

As observed for *E. coli* termination sequences, Cytosines are enriched over Guanosine adjacent to the TTS in *P. furiosus* (Fig. 2e). Motifs detected in the *E. coli* data set correspond to intrinsic (poly(U)) termination signatures and REP sequences^74,75^ that can frequently be found in intergenic regions and cause transcription termination at Rho-dependent attenuators (see Supplementary Fig. 7a,b)^28^. As expected, fold stability analysis of intrinsic *E. coli* terminators and hairpin-forming REP sequences, revealed secondary structures in both cases (Supplementary Fig. 7e). However, the stem-loops could potentially also represent processing or pause sites.

### Annotation of large transcriptional units

Long-read sequencing of native full-length RNAs has the potential to improve and facilitate genome-wide transcriptional unit (TU) annotation, which can be visually explored in a genome browser coverage track (Fig. 3a).

For whole-genome analysis, the annotation strategy was based on two major observations: First, during RNA preparation, RNA processing or degradation can occur, which limits the probability of sequencing an RNA in its native form as the percentage of full-length transcripts decreases with expected gene size (see Supplementary Fig. 9a). Secondly, we detected a decrease in coverage from the 3’ to 5’end of the RNA in all RNA classes except for the spike-in control (see Supplementary Fig. 9b), which is a limitation reported in the literature^13,76,77^. Therefore, we assume that not Nanopore sequencing but library preparation causes this problem. Based on this information, we developed a strategy that first collapses all overlapping reads and then splits them according to a significant coverage drop on the 3’ ends (annotation of TUs based on this strategy in Supplementary Table 5). We compared the results to database annotations and found that most of the differences are either caused by the low sequencing depth or by single-unit operons that have been collapsed and are now two-unit operons in the ONT data sets (see Supplementary Fig. 10a,b)^31,60^. Even though limited read availability is a concern in all data sets, many large operons were detected for all organisms (see Supplementary Fig. 10c). In case of limited bioinformatical resources, TUs can be explored visually in a genome browser, which is mostly not possible for Illumina reads (Fig. 3, see Supplementary Fig. 11,12). It further allows a quantitative analysis of individual transcripts in relation to other elements of the TU and performs much better than pure bioinformatical prediction or molecular biology methods (RT-PCR) as shown for the flagellum/archaellum operon in *P. furiosus* (Fig. 3)^60,78^. Here, it was possible to (i) detect multiple transcription units forming this cluster, (ii) confirm transcriptional start sites and (iii) to confirm that flaB0, the protein that is referred to as the major archaellin in *P. furiosus*^2,6^, is transcribed in large excess over the other archaellum genes. The largest identified TU cluster in *H. volcanii* mainly consists of ribosomal protein genes. Based on the native RNA-seq data, the analysis suggests that this operon is split into two transcription units. This shows that the ONT native RNA sequencing method provides the opportunity to annotate transcriptional units thereby outperforming the bioinformatics-only prediction as well as the visual inspection of Illumina coverage (see Supplementary Fig. 11). Besides, we confirmed the complex transcription pattern of the major ribosomal protein gene cluster in *E. coli* that stretches over more than 10 kb, including the accurate determination of TSS and TTS and a putative cleavage site in the secY gene (see Supplementary Fig. 12)^79^.

### Detection and confirmation of rRNA processing in *E. coli*

Next, we aimed to analyse the multi-step rRNA processing pathway which is the major RNA maturation pathway in any prokaryotic cell. We first focus on the *E. coli* data set as the processing of bacterial rRNAs is well characterized^80–82^. Ribosomal RNA in *E. coli* is transcribed from 7 independent rDNA operons encoding the mature rRNAs (16S, 23S and 5S rRNAs) and some tRNAs which are interspersed by RNA spacer elements^83^. In agreement with a previous study, transcription of *rrnC* from two promoters (transcription start sites at −293 and −175) was detected accurately in the TEX-treated sample, which is enriched in primary transcripts (Fig. 4a,b)^84^.

The rRNA maturation process, which requires the action of well-defined endo- and exo-ribonuclease activities, culminates in the formation of stoichiometric amounts of mature 16S, 23S, and 5S rRNAs^80–82,85^. Unexpectedly, the sequencing efficiency of mature 16S rRNA was lower than the 23S rRNA (Fig. 1b, see Supplementary Fig. 13). The reasons for this apparent discrepancy is so far unclear.

To re-trace the multi-step rRNA maturation process, we performed a co-occurrence analysis of read start and read end positions. Strikingly, we could identify most of the known 5’-processing/intermediate sites at nucleotide resolution in wildtype *E. coli* (Fig. 4b). Next, we categorized reads based on their experimentally verified and literature expected 5’ terminal positions and analysed 3’-enriched connected positions (Fig. 4c). Considering the 3’-to-5’ sequencing strategy of Nanopore sequencing, this co-occurrence analysis allows the assignment of 3’ terminal positions and distinction to random 3’ degraded reads.

Although we could detect RNA of similar size or longer (see above) very well, the short-lived full rDNA operon transcript detected in RNase III deficient strain^86^, is not observed using our experimental set-up. In contrast, the downstream known pre-rRNA intermediates, which are generated by the action of RNAse III were detected (Fig.4). Among these intermediates, the 17S pre-rRNA (115 additional nt at the 5’ end and 33 nt at the 3’ end of the 16S rRNA) and the P23S (7 additional nt at the 5’ end and 8 nt at the 3’ end of the 23S rRNA), were identified (Fig. 4b,c,d). Final 5’ end maturation of the 16S rRNA mainly occurs before the 3’ end^80^ by the action of additional ribonucleases (RNase E −66, RNase G −3, RNase AM 0 5’ mature^85^), which leads to an enrichment of reads that have extended 3’trailing regions compared to the mature position (Fig. 4c,d). Together we could identify most of the known rRNA processing-intermediates/-sites at near-nucleotide resolution in wildtype *E. coli*. However, it should be noted that the current experimental set-up can be biased by 5’ and 3’ degradation events, prohibiting precise 3’ end mapping in some cases and causing difficulties to identify short-lived/low-aboundant pre-rRNA intermediates.

### Insights into archaeal ribosomal RNA processing

In comparison to bacteria or eukaryotes, ribosomal RNA processing in archaea is still poorly understood^87–90^. Our current knowledge suggests that the primary polycistronic rRNA precursor contains two processing stems formed by the 5’ leader and 3’ trailer sequences surrounding the 16S and 23S rRNAs^88,89,91,92^. In Euryarchaeota, the 16S and 23S rRNAs are additionally separated by the presence of an internal tRNA. In most archaea, the 16S and 23S rRNA processing stems contain a bulge-helix-bulge (bhb) motif which is, in the context of intron-containing tRNA, recognized by the splicing endonuclease endA^91–93^. Similar to intron-containing tRNA maturation, processing at the bulge-helix-bulge motifs is followed by the covalent ligation of the resulting extremities, thereby generating the archaeal specific circular pre-16S and circular pre-23S rRNAs^89,92,94,95^. The exact molecular mechanisms by which the circular pre-rRNA intermediates are further processed into linear mature rRNAs remain to be fully characterized ^89,92,96^.

Performing enrichment analysis of terminal positions, we aimed to confirm and expand our knowledge on the poorly characterized multi-step ribosomal maturation process in two evolutionary divergent archaea, *P. furiosus* and *H. volcanii* (Fig. 5, see Supplementary Fig. 14)^58,89,91,97^. As expected, almost all reads are categorized as fully maturated transcripts of the single 16S/23S rRNA cluster that do not contain extended 5’ or 3’ spacer regions (Fig. 5c). Surprisingly, and in contrast to our analysis performed in *E. coli* (Fig. 4), some of the observed mature rRNAs 5’ positions did not precisely match the available annotations at NCBI (https://www.ncbi.nlm.nih.gov/genome/) or the archaeal genome browser (AGB, http://archaea.ucsc.edu), which are also showing discrepancies (summarized in Supplementary Figure 14). However, selected examination of the putative mature rRNA extremities obtained by ONT did match our independent experimental validations by primer extension analysis of the 5’ends of the 16S and 23S rRNAs of *H. volcanii* (Supplementary Fig. 14d). These results, and those obtained for *E. coli*, suggest that the mature 5’ of the rRNAs determined by native RNA sequencing most probably represent the genuine mature rRNA extremities.

Despite the high sequencing depth of the (pre-)rRNA, we did not detect a full-length precursor consisting of the 16S leading-16S-tRNA-23S-23S trailing elements in *P. furiosus* and *H. volcanii*, suggesting that, similar to the *E. coli* situation (see above), very early rRNA processing events may occur rapidly in these cells. The remaining rRNA reads were grouped according to (i) their 5’ leading and 3’ trailing lengths, (ii) the number of junctions and (iii) clipping properties of the alignments into several additional categories that are overall less abundant than the mature rRNAs and may represent either rRNA processing intermediates or are RNA elements generated as a product of pre-rRNA processing. Among these putative pre-rRNA-related intermediates, some are common to both archaea analysed, whereas others are apparently only found in one or the other organism. The pre-rRNA-related intermediates were selected on the basis of abundance and/or biological interpretability and/or prior characterization. The overall findings were used to extract an hypothetical rRNA maturation pathway in archaea which is summarized in Figure 5a. The rational for the selected pre-rRNA intermediates for *H. volcanii* and *P. furiosus* is described in more detail below and examplified in Figure 5b.

In *H. volcanii*, we identified 3 classes of putative intermediates. In the first class (class P2) the pre-rRNA boundaries of these intermediates match the previously described bhb processing sites located within the 16S and 23S rRNA processing stems, respectively (see Supplementary Fig. 14). These intermediates extend from the 5’ to 3’bulge cleavage sites, however, these extremities are not covalently ligated and may correspond to post-bhb cleavage/pre-ligation pre-rRNA intermediates. An exemplary verification of the 5’ boundary of the post-bhb cleavage/pre-ligation pre-23S rRNA analysed by primer extension is provided in Supplementary Figure 14d. The second class (class C) correspond to permuted reads covalently connecting the 5’ and 3’ bulge cleavage sites, and are likely observed as the result of random nicking of the circular pre-rRNA intermediates during sample preparations and are categorized as post-bhb/post-ligation pre-rRNA intermediates. We verified reads by re-mapping Nanopore and Illumina reads to a permuted RNA sequence that was designed by joining the 3’ bulge with the 5’ bulge to mimic the actual sequence of circular rRNAs (Fig. 5, see Supplementary Fig. 15). Similarly, we detected a third main class (class O), which corresponds to a putative pre-16S rRNA intermediate showing an immature 3’ end, which is extended by the typical permuted spacers sequence observed in the circular-pre-16S rRNA^95^. This topology possibly results from linearization of the circular pre-16S rRNA intermediate at the mature 16S rRNA 5’end (opened-circular-pre-16S rRNA). This putative pre-rRNA intermediate is relatively abundant in *H. volcanii* (n: 1120, 15% of all reads mapping in the 16S rRNA region) and strikingly shows a non random 3’end extremity - matching with the linearization of circ-pre-rRNA at the mature 16S rRNA 5’end. In contrast, the resulting 5’end were rather heterogenous, probably due to degradation during sample preparation (see Supplementary Fig. 15b).

To provide additional examples that show the potential to describe rRNA processing events in archaea, we sequenced an *H. volcanii* wildtype strain grown under low salt conditions known to accumulate large amounts of a longer 16S rRNA variant (see Supplementary Fig. 14b,c)^30^. Under these conditions, a 16S rRNA (precursor) with extended 5’ and 3’ UTRs (5’: −108 5’bulge, 3’: +70 3’bulge) appears that is enriched in this context. Quantification by comparison with the NOTEX wildtype set confirms the previous detection of this rRNA variant in a gel-electrophoretic analysis of total RNA (see Supplementary Fig. 14b)^30^. To reveal more details about the nature of these precursors, we re-mapped the reads to the permuted linear rRNA 16S sequence. We observed that the relative number of reads obtained for circ-pre-16S rRNA (class C) and especially of opened-circ-pre-16S rRNA (class O) obtained under low salt conditions were exceeding the ones from “normal” conditions, indicating that rRNA maturation and/or turnover is affected in this “stress” condition (Fig. 5d). The functional relevance of these observations remain to be analysed.

In *P. furiosus*, for which we have obtained larger amounts of reads, we could define 4 categories of pre-rRNA-related intermediates (see Supplementary Fig. 14k, ranked by their timely appearance): (1) Fragmented full-length precursor rRNAs (I), (2) 16S rRNA leading/trailing sequence-tRNA-23S rRNA, (3) a putative RNA chimera resulting from processing and RNA ligation activities that entails the 16S rRNA leading/trailing sequence-tRNA-23S rRNA leading/trailing sequence (T1), and (4) permuted 16S and 23S pre-rRNA intermediates (C) (Fig. 5, see Supplementary Fig. 14).

The putative permuted reads (4) are reminiscent of the reads typically observed for circular pre-rRNA in *H. volcanii* (class C) and may correspond to the covalent ligation of the 5’ and 3’ spacers generated by cleavages at the bulge-helix-bulge motifs within the processing stems (Figure 5 and ^92,94,95^). To verify this hypothesis, we performed RNA structure prediction of the corresponding double stranded RNA regions (see Supplementary Fig. 16). In agreement with the permuted reads, we could place the corresponding extremities within the bulge-helix-bulge motifs. However, the 23S processing stem does not adopt a canonical bhb motif, but forms an alternative structure similar to the one previously described for the 16S rRNA bhb motif in *S. acidocaldarius*^93,98^. Others and we could previously demonstrate that this alternative structure is compatible with circular-pre-16S rRNA formation in *S. acidocaldarius*^94,95^. Therefore, these permuted reads likely originate from random opening of the archaeal specific circular-pre-rRNA intermediates during sample preparation/sequencing (as observed for *H. volcanii*) and suggest that like various archaea analysed so far, circular pre-rRNA intermediates are also produced in *P. furiosus*. Although we could detect a similar total number of circular pre-rRNA transcripts for *H. volcanii* and *P. furiosus*, the proportion of this category with respect to all reads mapping to the 16S rRNA is very low, which might reflect the actual abundance (Fig. 5e). However, considering the differing sequencing efficiencies of the 16S rRNAs absolute quantifications cannot be made.

The early RNA chimera 16S rRNA leading/trailing sequence-tRNA-23S rRNA trailing sequence precursor (T1) likely generated by cleavage and reciprocal ligation of the pre-16S and pre-23S rRNAs at the predicted bulge-helix-bulge motifs were detected very accurately, and are reminiscent of previous observations^92^ (Fig. 5, see Supplementary Fig. 14 l). Given the number of reads, the direction of ONT sequencing from 3’ to 5’ and the accurate mapping, it is unlikely that the additional putative rRNA precursor (P1) carrying the leading sequence in combination with tRNA-23S rRNA is arising from an experimental artifact (Fig. 5, see Supplementary Fig. 14). In fact, this variant is in good agreement with our recent *cis*-acting element analysis in *H. volcanii*^95^ (see also Discussion).

Taken together, our analysis confirms and expands the number of putative pre-rRNA intermediates in archaea. Moreover, this extended framework provides an additional basis to facilitate further definition of common and specific principles of rRNA maturation in archaea.

### Towards mapping of RNA base modifications

More than 160 types of modified bases have been described in RNAs so far^99^. In contrast to other sequencing techniques, Nanopore-based sequencing offers the possibility to detect base modifications directly as these modifications lead to an electric current signal that differs from the expected theoretical distribution obtained by the unmodified nucleotide sequence^15,16,20,100^. In turn, these signal changes might also lead to differences in the basecalling profiles (e.g. systematic errors or a drop in basecalling quality). Approaches based on signal deviations or basecalling errors have already been applied to map RNA and DNA modifications in different organisms. However, accurate *de novo* RNA modification prediction with single-nucleotide resolution is still challenging as more than one base affects the current through the pore (see Fig. 6a). In addition, current deviation is influenced by the type of modification and the surrounding sequence context^101^.

Despite these limitations, we aimed to study rRNA modifications in archaea and explored different analysis strategies (Fig. 6). Based on the approaches mentioned above, we benchmarked the potential to detect known and putative modification sites in the 16S rRNA. We focused first on the 16S rRNA modifications of *P. furiosus*, using the recently established 16S rRNA modification pattern in the close relative *Pyrococcus abyssi*^59^. This set includes 34 N^4^-acetylcytidines, and 10 other modifications of diverse types^59^. Compared to a background set consisting of all other positions in the *P. furiosus* 16S rRNA, we observed that the surrounding sequencing context of all modified bases is significantly enriched in basecalling errors (Fig. 6b) and also had a comparatively low mapping quality (Fig. 6c). Depending on the type of modification (acetylation vs. diverse), these metrics looked very different across a sequence context from −5 to +5 from the exact position of the modified base (Fig. 6d, see Supplementary Fig. 17a,b). While N^4^-acetylcytidines were mostly miscalled at their predicted position, the other diverse base modifications had various effects on all metrics (Fig. 6b, see Supplementary Fig. 17a,b). Using the position information derived from *P. abyssi*, our analysis suggest that the putative N^4^-acetylcytidine modification leads to a wrong base assignment during basecalling (Fig. 6d,e). In fact, this systematic non-random error was also reflected by the high proportion of a central T instead of C in the CCG context of the acetlyation (see Supplementary Fig. 17c,d).

To analyse whether these modifications are already established at early steps of rRNA maturation, we looked at the basecalling properties of 5’extended pre-rRNAs and compared them to mature 16S rRNA. Importantly, we did not observe significant basecalling errors in these selected precursor rRNAs (5’extended pre-16S RNA) indicating that the cytidine N^4^-acetylation is not occurring early in the rRNA maturation pathway (see Supplementary Fig. 17e) (see below for further details).

As the approach based on systematic errors gave us promising results for N^4^-acetylations, but were less unambiguous for diverse modifications, we wanted to evaluate the potential to detect RNA base modification from raw signals using Tombo (Fig. 6a, updated Tombo version from Stoiber et al.^57^). To this end, we first focused on the dimethylation (m^6^_2_A) introduced by the enzyme KsgA/Dim1 at position A1450/A1451 in *H. volcanii* (A1518/A1519 *E. coli* numbering)^58,102^. Using the *de novo* model in Tombo the calculated probability of a modification was very high for the stretch of Guanosines adjacent to position A1450 (Fig. 6f). Mapping to single-nucleotide resolution is difficult as more than one base contributes to the actual electric current signal in the nanopore^103^. In the next step, a comparison of a wildtype sample to a deletion mutant of archaeal KsgA/Dim1 homologue helped us to confirm that the current signal alteration in this particular region is dependent on the KsgA/Dim1 m^6^_2_A modifications and not the homopolymer-G-stretch (Fig. 6g). The analysis further revealed a reduced signal variability at non-modified positions between the two samples in comparison to the theoretical distribution, which leads to less false positives in the statistical analysis and highlights the benefits of a background read model.

Assuming that early pre-rRNAs represent a state where base modifications are not yet quantitatively introduced, we used these reads as a background model to explore the potential to detect the introduction of base modifications at different stages of rRNA maturation in archaea. Therefore, we generated multiple sets by sorting reads according to the main classes of pre-rRNA intermediates described above for *H. volcanii* and *P. furiosus* (Fig. 7a). For *H. volcanii* and *P. furiosus*, we compared 5’extended 16S rRNA, circular pre-16S rRNA, opened circular-pre-16S rRNA and mature 16S rRNA. We first focussed on analysing the m^6^_2_A and N^4^-acetylcytidine signatures across the putative different stage of rRNA maturation in archaea. To this end, we evaluated basecalling and raw signal profiles using the 5’ extended pre-rRNA as a background model as the former performed well for N^4^-acetylcytidines, while the latter for m^6^_2_A detection.

Basecalling anomaly in the m^6^_2_A region was detected within the mature 16S rRNA of both *P. furiosus* and *H. volcanii*, and could be confirmed by a high probability of base modification around this positions using the tombo model (Fig. 7b,c, see Supplementary Fig. 18). Interestingly, similar profiles were detected for the putative opened-circ-pre-16S rRNA in *H. volcanii*, but not in *P. furiosus*. This finding is in line with a proposed “quality control” function of KsgA/Dim1 during late biogenesis of the small ribosomal subunit^104–106^. Similarly, we analysed N^4^-acetylcytidine known to occur in the vicinity of the m^6^_2_A. However, this modification occurs prior to the KsgA/Dim1-dependent modification during eukaryotic ribosome biogenesis^107–111^. In agreement with previous analysis^58^, no apparent N^4^-acetylcytidine modification was observed at the equivalent position in *H. volcanii* (Fig. 7c). In contrast, an increase in base-calling errors at the expected position was observed in circular pre-rRNAs and mature 16S rRNA, but not within 5’extended pre-16S rRNA in *P. furiosus*. Therefore, and similarly in eukaryotes, N^4^-acetylcytidine modification in helix 45 precede the KsgA/Dim1-dependent m^6^_2_A modifications in *P. furiosus*. Note that the results for the circular pre-16S rRNA (grey, Fig. 7) should be taken with care in *H. volcanii*, given the limited number of reads in this category (see Supplementary Fig. 18). However, it is tempting to speculate that a different timing of events in *H. volcanii*, observed by the earlier m^6^_2_A modifications, is caused by the absence of N^4^-acetylations.

To further extend on the timely order of all N^4^-acetylcytidine modification potentially present in *P. furiosus*, we analysed the basecalling profiles across different rRNA maturation stages (Fig. 7d). In addition, we compared it to all other CCGs that are present in 16S and 23S rRNA as N^4^-acetylation have been shown to be introduced in a CCG context in *P. abyssi*^*59*^. This analysis suggests that (i) N^4^-acetylcytidine modifications may be also scattered across the 23S rRNA sequence, and (ii) these modifications are established in the course of pre-rRNA maturation (circ-pre-rRNA in Figure 7d). Taken together, our analysis, suggests that despite the current limitations, ONT allows to discriminate (some) rRNA modifications across selected rRNA maturation events. Moreover, these data support the long-standing hypothesis that hyperthermophilic organisms might stabilize their rRNAs by a higher degree of RNA modifications^112,113^.

## Discussion

Performing whole-transcriptome native RNA-seq study in prokaryotes provided us with a wealth of information on transcriptional and post-transcriptional processes in *E. coli* and the archaeal model organisms *H. volcanii* and *P. furiosus*. Here, we will mostly discuss new biological insights that emerged from our study. Additionally, we will reflect on the advantages and disadvantages of Nanopore native RNA-seq.

### Insights into transcriptional processes

Bacterial and archaeal transcription is an intensely studied molecular process and the mechanisms of basal transcription are well understood^114^. Native RNA sequencing allowed us to retrieve accurate information of transcript boundaries on both 5’ and 3’ ends. Our data show that 3’ UTRs length distributions are comparable between *E. coli, P. furiosus* and *H. volcanii* with the majority of mRNAs showing a length between 30-70 nt. Similar to bacteria, archaea encode a large number of small non-coding RNAs^115^. However, many regulatory events that involve the regulation via small RNAs take place at bacterial 5’ UTRs^116^. We and others found that 5’ UTRs are significantly shorter in many archaea supporting the idea that post-transcriptional regulation is mediated via the 3’ rather than the 5’ UTR in these groups^117^. Additionally, we determined transcription termination sites, which are well analysed for bacterial species but only a few studies focused on archaeal termination mechanisms, especially on the genome-wide level. In both archaeal species studied, poly(U) stretches were overrepresented at termination sites agreeing well with termination sequences found in *Sulfolobus* and *Methanosarcina*^54^. Interestingly, the majority of TTS found in *Pyrococcus* transcripts is composed of two U-stretches with at least five consecutive uridine bases while a subclass of *Haloferax* transcripts is almost exclusively terminated by a single U-stretch with four uridine bases. It has been shown that a five base U-stretch is sufficient to induce termination *in vitro*^72,118,119^. Similar observations were described in a recent study by Berkemer et al, which identified a poly(U)_4_ stretch to be the termination signal in intergenic regions^56,59^. Notably, the *H. volcanii* genome is distinguised by a high GC content leading to a low propability for the occurence of U_5_ stretches and hence, the transcription machinery might have adapted to recognise U_4_ stretches as termination signal. However, the current data set suggests that this short termination signal might be a specific feature for a subclass of *Haloferax* transcripts resembling the poly(U) termination motif found in *E. coli*. All other archaeal organisms (*P. furiosus, M. mazei, S. acidocaldarius*) investigated so far terminate transcription at multiple consecutive poly(U) stretches. Possibly, *Haloferax* relies on additional termination signals or yet unknown termination factors. A putative candidate is archaeal CPSF1 (aCPSF1, also known as FttA), a recently described archaeal termination factor^70,120^ that is widespread in archaea. aCPSF1 acts as ribonuclease that was shown to cleave transcripts after a poly(U) stretch to trim transcripts and facilitates transcription termination in *Thermococcus kodakarensis*^70^ and *Methanococcus maripaludis*^120^. The arising 3’ UTR isoforms were detected using Term-seq analysis^120^. We also observed heterogeneity in the case of the Pilin and histone transcripts, respectively, that are distinguished by varying lengths of the 3’ UTR suggesting that aCPSF1 might trim a subset of genes in *H. volcanii* and *P. furiosus*. It is noteworthy that 3’ UTR isoforms were also detected in Term-seq studies with *Sulfolobus* and *Methanosarcina*^54^. However, in contrast to the Pilin, Alba and histone transcripts, the 3’ UTR isoforms arise from termination at different sites of a single continues poly(U) stretch suggesting that the isoforms arise from a stochastic termination process of the RNA polymerases at an extended poly(U) stretch at the end of the gene. The gradual termination observed in this study might also be influenced by the coupling of transcription and translation. These genes are all highly expressed and translated. Hence, it seems plausible that the ribosome is efficiently coupled to the RNAP^121^ (as observed in bacteria^38,38,122,123^). Several studies in bacteria showed that the ribosome influences transcription (and *vice versa*)^124–126^. The stochastic termination might therefore be a result of the uncoupling of the ribosome at the end of the mRNA potentially also inducing the dissociation of the transcription elongation complex. Taken together, these data suggest that a variety of termination mechanisms (that can even co-occur in the same cell) can be found in archaea ranging from stochastic intrinsic termination at extended poly(U) stretches (*Pyrococcus, Sulfolobus, Methanosarcina*), to abrupt termination at short uridine tracts (*H. volcanii*) and factor-dependent termination that results in trimming of the 3’UTR (*H. volcanii, P. furiosus, M. maripaludis, T. kodakarensis*).

In the context of transcription, the long and overlapping native RNA reads helped us to analyse the transcriptional landscape at multigene operons. More specifically, we focused on the archaeal flaggelum (archaellum) operon, encoding for the archaeal motility machinery^127^, as the transcription unit assignment remained ambiguous so far^78^. In contrast to bioinformatical and Illumina RNA-seq-based predictions and attempts to unravel the TU via primer extension experiments, we found that the archaellum operon in *P. furiosus* is transcribed in multiple units with highly diverse abundances. The *fla*B0 gene encodes the major archaellin/flagellin protein that forms the filament of the archaellum and therefore, the organism has to produce this protein in large quantities as apparent from the expression level^78^. Interestingly, FlaD mRNA is expressed at comparably high levels as well supporting the idea that FlaD is a major constituent of the archaellum in *P. furiosus*. It has been speculated that FlaD forms the cytosolic ring of the archaellum that anchors the filament in the outer membrane^128^. The identity and functional role of FlaD are, however, not known so far.

### Insights into rRNA processing in archaea

In this study, we have assessed the suitability of native RNA sequencing to obtain information on the rRNA maturation pathway of different prokaryotes. Ribosomal RNA processing proceeds via the coordinated and defined order of ribonucleases action (exonucleolytic and/or endonucleolytic cleavages) which generate pre-rRNA intermediates with defined premature rRNA sequences^82,90,129,130^. The establishment of faithfull rRNA maturation maps in model organisms, like *E. coli, S. cerevisiae* or human cell culture has required numerous analyses over the past decades^82,90,129,130^, and remains a technical challenge. Therefore, methodologies that might accelerate the systematic analysis of rRNA maturation pathways across the tree of life, thereby enabling to unravel the diversity of rRNA processing strategies need to be established. Beyond the identification of processing sites, the order of the processing events which can be, in part, deduced from co-occurrence analysis of the 5’ and 3’ extremities is of biological relevance^82,90,129,130^. Whereas we could confirm and extend our general view on the rRNA maturation pathway in archaea, the 3’-5’ processivity of Nanopore native RNA sequencing observed for rRNA and the potential RNA degradation during sample preparation impedes the accurate quantitative analysis of pre-rRNA extremities co-segregation (see Fig. 5 and Supplementary Fig. 9b). Nevertheless, we could, in most of the cases, confirm and expand the presence of pre-rRNA intermediates and processing sites in the different organisms analysed, including the archaeal specific circular-pre-rRNA intermediates^89,92,94–96^(see discussion below). Together our findings are summarized into an updated archaeal rRNA processing model described in figure 5 and are discussed below.

The full length theoretical primary rRNA transcript was not identified in any of the archaeal organisms analysed. Similarly, this primary rRNA is generally difficult to observe in wildtype *E. coli* (^86,131^ and this work). Collectively, these observations suggest that short-lived and/or low abundant pre-rRNA intermediates escape the detection capacity of the current experimental set-up. Accordingly, it is also difficult to infer differences in rRNA processing features between different (archaeal) organisms by virtue of observed pre-rRNA intermediates absence/presence pattern. In fact, these differences may also be related to organism-specific changes in pre-rRNA intermediates relative levels, which will depend on the sum of the reaction kinetics of the different maturation steps in a given condition.

Among the identified pre-rRNA intermediates, the *post*-16S-bhb/*pre*-ligation precursor (P1), which is observed in *P. furiosus* and includes ligation at the bhb motif of the upstream region of the 16S leader and downstream region of the 16S trailer sequences and continues to the downstream tRNA/23S sequences, is of particular interest (see Supplementary Fig. 14k). The presence of this ligation event suggests that the 16S rRNA bulge-helix-bulge processing occurs prior to internal tRNA and 23S rRNA maturation. Although, this ligation event was not identified by ONT in *H. volcanii*, this observation is in agreement with our recent functional *cis*-acting element analysis performed in *H. volcanii*^89,92,94^. In fact, based on this previous analysis we have proposed a model by which 16S rRNA maturation proceeds and is required for the downstream maturation of the internal tRNA and 23S rRNA. Moreover, we have hypothesized that ligation of the 16S rRNA leader/trailer resulting from the 16S rRNA bulge-helix-bulge maturation process generates a putative new pre-rRNA intermediate for which the corresponding ligation event could be observed in *Pyrococcus furiosus* using native RNA sequencing^95^. In addition, the presence of an RNA chimera containing the leading/trailing/tRNA parts (T1) (*post*-bhb/*post*-ligation) support the idea that the maturation of the co-transcribed internal tRNA is inefficient or inhibited and may preferentially occur after processing of the 16S and 23S rRNA bulge-helix-bulge which liberate the circular pre-16S and pre-23S rRNAs (C) (suggested in^92,95^, and this work). The presence of circular pre-16S and pre-23S rRNAs and their processing sites could be verified and established in *H. volcanii* and *P. furiosus*, respectively (Fig. 5, see Supplementary Fig. 15/16). Recently, we determined the functional requirement of the bulge-helix-bulge motifs for the formation of circ-pre-rRNAs in *H. volcanii*. Moreover, in analogy to intron containing-tRNA splicing, the rRNA bhb motifs are presumably cleaved by the tRNA splicing endonuclease (endA) prior to covalent circularization^91,95,96^. Although intact circular RNA can not be directly sequenced by ONT, we noticed the presence of permuted transcript in *H. volcanii* corresponding to the ligation events previously identified for circ-pre-rRNAs in *H. volcanii*^*95*^. Most of these permuted reads were also showing random and heterogenous 5’and 3’ends thereby suggesting that these pre-rRNAs were likely the result of randomly nicking of circular pre-rRNA intermediates during sample preparation (see Supplementary Fig. 15). Noteworthy, similar permuted reads were observed in *P. furiosus*, for which the presence of circular-pre-rRNA intermediates is not established thus far. Whereas, the observed ligation could be accurately mapped to the predicted 16S bhb motif, the 23S bhb motif could not be accurately predicted (data not shown). However, our manual inspection suggests that the permuted reads extremities match to an imperfect, presumably less stable, bhb motif within the 23S processing stem (see Supplemantary Fig. 16b). This property is reminiscent to the “aberrant” 16S bhb motif used for circular-pre-16S rRNA formation in *S. acidocaldarius*^*93–95,98*^. Whether these structural features are stabilized by additional factors or enable a certain degree of regulation during the rRNA maturation process in the cellular context is unknown.

In addition to the circular pre-rRNAs, we observed pre-rRNA intermediates cleaved at the bhb motifs but not yet ligated into circular pre-rRNA in *H. volcanii (post-*bhb*/pre-*ligation pre-rRNAs) (P2). Whereas, the presence of this intermediate processing step is theoretically expected, they were only detectable in *H. volcanii* (see Fig. 5a, see Supplementary Fig. 14), suggesting that the maturation kinetics or stability of these pre-rRNA intermediates varies among these organisms. How the circular pre-rRNAs are further processed into linear mature rRNA is not well understood. Based on our current knowledge, several non-mutually exclusive hypothesis can be drawn: (i) opening of the circular-pre-rRNA within the ligated spacer region and subsequent maturation of the 5’and 3’end; (ii) opening of the circular pre-rRNA by first maturation of the 5’end mature rRNA followed by 3’end maturation; or (iii) opening of the circular pre-rRNA by first maturation of the 3’end mature rRNA followed by 5’end maturation. A category of putative 16S pre-rRNA intermediates observed in *H. volcanii*, may provide some indications how linearization of the circular pre-16S rRNA is achieved. In fact, this particular intermediate was extended in its 3’end by the presence of the ligated 5’and 3’spacers normally observed in the circular pre-16S rRNAs and this 3’extension consistently ended just prior to the 16S 5’mature ends. This particular configuration is suggestive of 5’end maturation of circular-pre-16S rRNA prior to final 3’end maturation, thereby generating opened-circular pre-16S intermediates (O) (see Supplementary 15b). Although the majority of opened-circular pre-16S rRNAs is degraded from its 5’end, we detected a subset representing the theoretical full length (see Supplementary Fig. 15b,c). Additional properties of this putative intermediate is in agreement with its positioning during rRNA maturation (see below discussion on rRNA modifications) and with the prevalence of 16S rRNA 5’maturation prior to its 3’end previously observed in bacteria and eukaryotes^82,90^. Overall, future functional charaterization of the *cis*- and *trans*-acting elements required for pre-rRNAs maturation will be necessary to further refine our view on archaeal rRNA processing.

In conclusion, despite some intrinsic limitations, we provide evidence that direct RNA sequencing technologies can be a useful tool to approach intricated maturation pathway like rRNA maturation, and expand our understanding of RNA maturation in prokaryotes.

### Towards the mapping of rRNA modification patterns

RNA modifications have been described already in the 50-60s, and have gained significant attention over the last years, under the generic term of the epitranscriptome^132–134^. The high-throughput analysis of these post-transcriptional modifications remains challenging and mostly relies on indirect methods, like primer extension stops analysis and/or chemical recoding/derivation strategies^135,136^. Native RNA sequencing may fill an important gap to systematically analyse RNA modifications on a genome-wide scale. However, global strategies enabling the faithful determination of RNA modification identity and position needs to be developed. Several recent analyses have explored different strategies to evaluate the capacity of ONT to accurately detect RNA modifications (e.g. m^6^A)^15,16,65,137,138^.

RNA modifications can lead to electric current signals varying from the theoretical signal expected for the unmodified canonical ribonucleotides. These properties can be harnessed, on the one hand, to predict RNA modification probability by comparing theoretical and experimental electric current signal distribution, and on the other hand, variation of electric signals may increase the rate of basecalling errors. In both cases, the comparison of the profiles to a background distribution of non-modified nucleotides leads to a significant reduction of false-positives. We evaluated the potential to use early rRNA precursors, which are expected to contain incomplete modification patterns, as a background model and applied this strategy to analyse different stages of rRNA maturation.

To validate our approach, we first focussed on two types of modification occurring in close proximity in helix 45 of the 16S/18S rRNA, but at distinct stages of rRNA maturation, namely the almost universally conserved KsgA-dependent dimethylations (m^6^_2_A) and the less conserved Kre33/Nat10-dependent N^4^-cytidine acetylation^16,102,105,139,140^. By analysing basecalling profiles and raw signals in wt and KsgA deletion strain we could unambiguously provide *in vivo* evidence that the archaeal KsgA-dependent methylations of the 16S rRNA are completed at a late stage of the small ribosomal subunit biogenesis in both *H. volcanii* and *P. furiosus*, and may predominantly occur after linearization of the circular-pre-16S rRNA. In contrast, helix 45 N^4^-cytidine acetylation, which is absent in *H. volcanii*, appears to be added at the circular-pre-16S rRNA stage, prior to completion of the KsgA-dependent modifications in *P. furiosus* (Fig. 7). These results are in good agreement with previous studies done in eukaryotes and bacteria^104–106,108–111,141^. Moreover, expanding our sample-compare approach also suggests an increased amount of rRNA modifications in the hyperthermophile *P. furiosus*, and a decrease amount of predicted rRNA modifications in halophile *H. volcanii* in comparison to *E. coli*. These differential modification patterns across archaea are in good agreement with previous studies and may reflect adaptation to the environmental conditions that these extremophilic archaea encounter^58,112,113^. Recently, it has been shown that *P. abyssi* 16S rRNA is heavily acetylated at CCG motifs^59^. Our analysis suggests that N^4^-acetylcytidine modifications (i) increases the rate of basecalling errors (e.g. C>T) at the expected modified residue, (ii) are distributed across the 16S and 23S rRNA sequences in *P. furiosus*, and (iii) are successively added during rRNA maturation to reach “completeness” in the mature rRNAs. Future studies will be necessary to decipher, how widespread this type of modification is among archaea, and to evaluate their contribution for ribosomal subunit biogenesis and function in the cellular context.

Whereas ONT may facilitate RNA modification analysis in general, the exact chemical nature of these modifications can not be unveiled without prior knowledge and remain a challenging task which greatly benefits of the use of unmodified/hypo-modified references (in agreement with recent studies^16,138^). To facilitate high-throughput identification of RNA (DNA) modifications, future studies will required to develop and train algorithms improving the *de novo* identification confidence of diverse RNA/DNA modifications.

### Benefits and limitations of Nanopore-based native RNA sequencing

Taken together, a key advantage of the native RNA-seq approach is that multiple features can be addressed at once distinguishing the technique from the Illumina sequencing technology or biochemical assays. ONT sequencing does not require large scale equipment and is a fast method. Moreover, the method does not necessitate a reverse transcription step or PCR amplification thereby avoiding biases introduced by these enzymes. Due to the limitations of the sequencing read analysis platform, ONT sequencing does not accurately detect small RNAs yet. Additional limitations of the native RNA-seq technique are currently (i) the high amount of input RNA required (2-5 µg) to reach good coverage of the transcriptome without rRNA depletion, (ii) the need for a enzymatic poly-adenylation step of non polyA+ RNA, (iii) the 3’ bias during RNA sequencing (iv) limited throughput and (v) limited possibilities for multiplexing. Although ONT sequencing has a comparably low sequencing accuracy, this did not pose a limitation for our analysis. Due to the extraordinary read length and the sensitivity to base modifications, ONT-based native RNA-seq can provide valuable insights into (r)RNA processing, (r)RNA modification patterns and the transcription of large operons. Strikingly, ONT-based sequencing is a *bona fide* single-molecule method and hence molecular heterogeneity in the transcriptome can be analysed so that even minor RNA populations can be detected that are inevitably lost in ensemble sequencing approaches.

## Data availability

Raw sequencing data sets (gzipped raw FAST5 files) will be deposited in the Sequence Read Archive (SRA) and will be available under project accession number PRJNA632538.

## Code availability

A detailed documentation and code of all essential analysis steps (used tools and custom Rscripts) are available from https://github.com/felixgrunberger/Native_RNAseq_Microbes.

## Author contributions

F.G. established the nanopore workflow and performed all the bioinformatic analysis. F.G., R.K., M.J., R.R. and A.B. performed RNA extractions. M.F. helped to optimize the RNA treatment protocol. F.G. carried out library preparations and performed sequencing. F.G., R.K., M.J. carried out *H. volcanii* wildtype/Δ*ksgA* library preparations and sequencing. M.F. and R.R. performed transcription assays. R.K. and S.F.-C. generated the KsgA deletion strain. R.K. performed primer extension analysis. F.G., S.F.-C. and D.G. designed the study, analysed and interpreted the data, and wrote the manuscript with the input of all authors. J.S., W.H., S.F.-C. and D.G. supervised the experiments. S.F.-C. and D.G. initiated and supervised the project.

## Acknowledgements

We gratefully acknowledge financial support by the Deutsche Forschungsgemeinschaft within the collaborative research center framework (CRC/SFB960) “RNP biogenesis: assembly of ribosomes and non-ribosomal RNPs and control of their function” [SFB960-TP7 to D.G.] [SFB960-TP-B13 to S.F.-C.]. The work was also supported by the DFG through grant So264/21 to J.S.

## Notes

https://github.com/felixgrunberger/Native_RNAseq_Microbes

### Competing Interest Statement

The authors have declared no competing interest.

### Summary of Updates

- additional and refined analysis of rRNA maturation pathway - additional analysis of map rRNA modifications - additional analysis to correlate rRNA modification to rRNA maturation steps

