## Supplementary Information for "Exploring prokaryotic transcription, operon structures, rRNA maturation and modifications using Nanopore-based native RNA sequencing"

Grünberger et al.

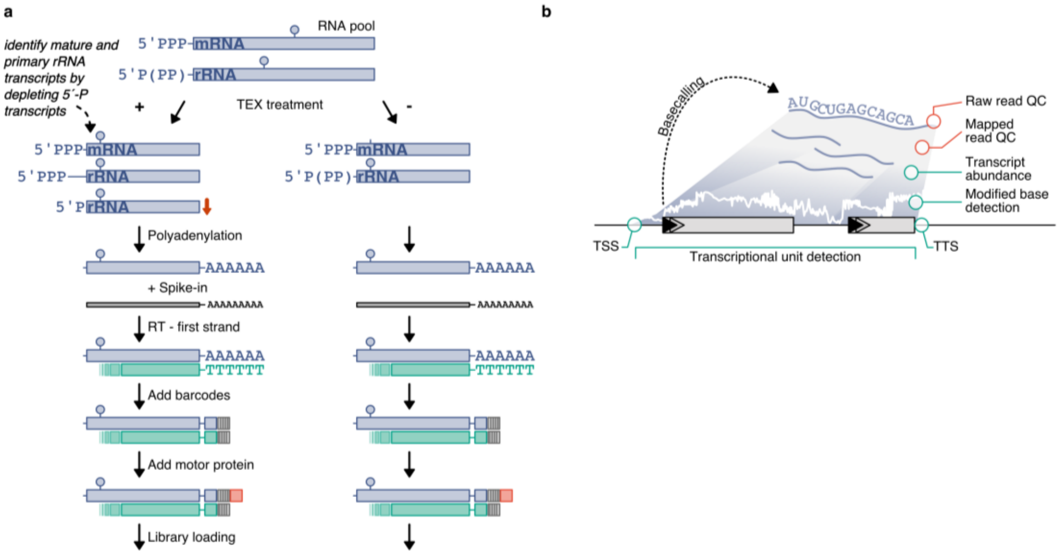


**Supplementary Figure 1 | Extended workflow and objectives of Nanopore-based native RNA sequencing. a**, Partial digestion of RNAs that are not 5´-triphosphorylated (e.g. tRNAs, rRNAs) is done using a Terminator 5´ -Phosphate-Dependent Exonuclease (TEX), allowing for the analysis of both mRNAs and rRNAs. After enzymatic poly(A)-addition, library preparation is performed according to a modified SQK-RNA001 protocol. The spike-in enolase is added and the RNA is reverse transcribed using the RTA adapter (green poly(T)). Depending on the necessary sequencing depth, custom barcodes (poreplex) can be added in the next step that replace the 3´- RMX adapter (lined grey square). A further 3´- ligation is performed to add the motor-protein carrying adapter (red square). Libraries are then loaded on a R9.4 flowcell and sequenced for 48 hours on a MinION device. **b**, After demultiplexing and basecalling of raw reads, quality control is performed on raw reads and mapped reads (red) to detect problems during library preparation or sequencing. Multiple transcriptomic features highlighted in green, including transcriptional start sites (TSS), termination sites (TTS), transcript abundance, modified bases and composition of transcriptional units can be adressed simultanously.

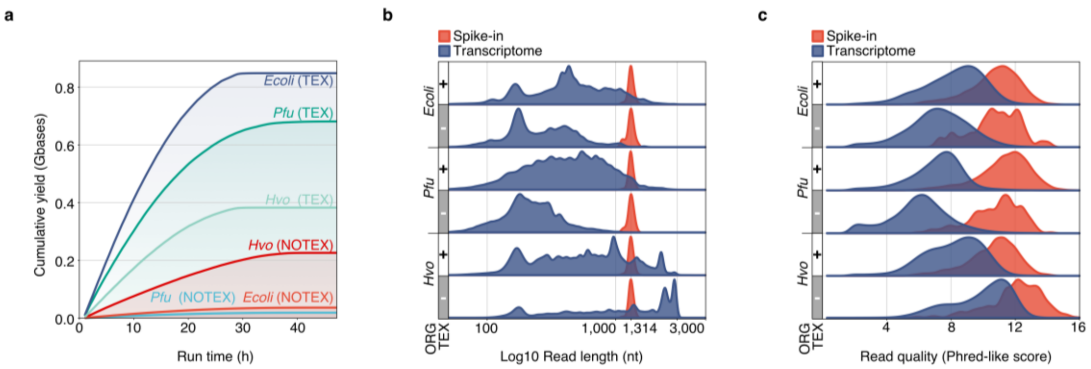


**Supplementary Figure 2 | Raw read analysis. a**, Libraries are loaded on R9.4 flow cells and reads are collected over 48 hours using the recommended MinKNOW script. FAST5 files are then demultiplexed (poreplex) and converted to FASTQ files during guppy basecalling. Differences in the plotted cumulative yield derive mostly from different multiplexing strategies. **b**, Raw read length ($log_{10}$ scale) and **c**, raw read quality (Phred-like score estimated by guppy) is compared between samples for transcriptome-derived reads (purple) and the spike-in control (red).

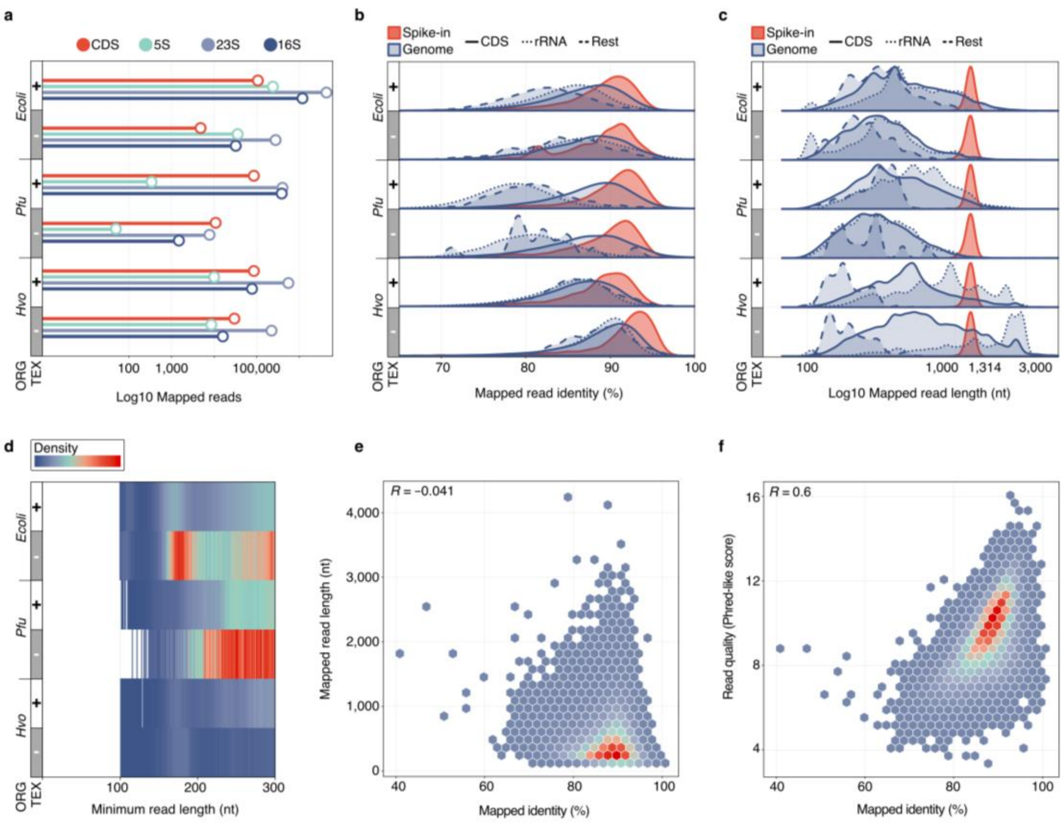


**Supplementary Figure 3 | Mapped read analysis. a**, Number of mapped reads ($Log_{10}$ scale) color-coded for different features (protein coding genes (CDS): red, 5S rRNA: green, 16S rRNA: purple, 23S rRNA: light-purple, compare Fig. 1b). **b**, Mapped read identity is defined as (1-NM/aligned_reads)*100, where NM is the edit distance reported taken from minimap2[1]. Distribution of read identity (in %) and **c**, mapped read length (in $Log_{10}$ scale) are shown for spike-in control (enolase yeast, red) and genomic features (purple). Different feature types are highlighted with solid line (CDS), dotted line (rRNA), long-dashed line (rest) and are based on the featurecounts classification[2]. **d**, Limitations in the minimum read length that can be mapped (in nts) is shown for all datasets. Density of reads is indicated by a color-scale from darkblue (few reads) to red (many reads). **e**, Mapped identity (%) is compared to mapped read lengths (nt) for the same sample (Spearman´s rho: -0.041). **f**, Raw read quality correlates with calculated mapped identity (Spearman´s rho = 0.6). Density is color-coded as described before. TEX treated *P. furiosus* set (CDS mapping reads) is shown as a representative example.

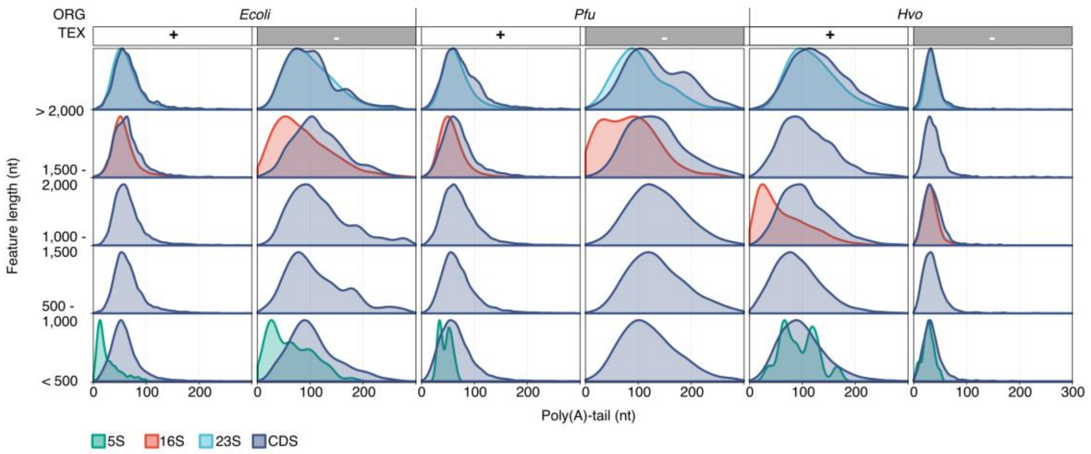


**Supplementary Figure 4 | Poly(A)-tailing efficiency.** Distribution of poly(A)-tails (nt) estimated by nanopolish[3] is shown for color-coded transcript groups that were separated based on their region lengths (protein coding genes (CDS): purple, 5S: green, 16S: red, 23S: blue).

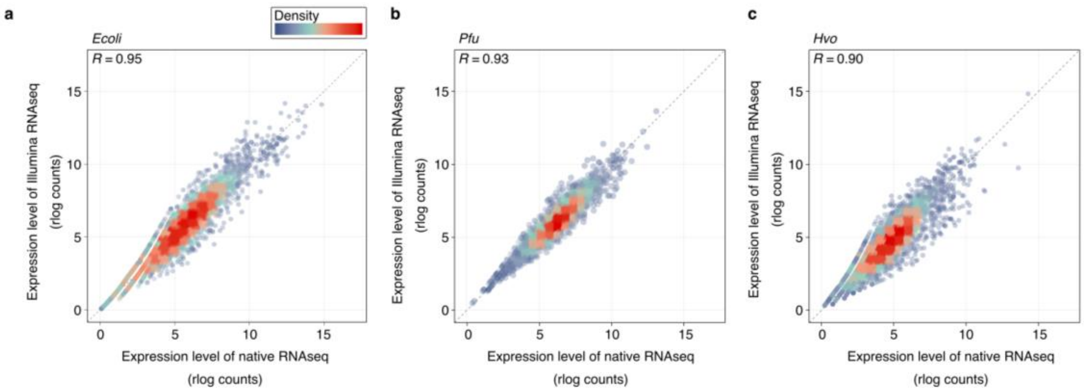


**Supplementary Figure 5 | Correlation of transcript abundance levels between Nanopore native RNA-Seq and Illumina RNA-Seq.** Transcript counts are calculated by featurecounts and rlog transformed using the DESeq2 package in R to account for differences in sample sizes[2,4]. Each gene is represented by a point colored by density from low (darkblue) to high (red) to address for overplotting. Correlation is calculated by Spearman´s rho and shown in the upper left. Publicly available data sets generated by Illumina sequencing (see detailed description in Material and Methods) were used to compare ONT samples in **a**, *E. coli* (SRP056485), **b**, *P. furiosus*[5] and **c**, *H. volcanii* (SRR7811297)[6].

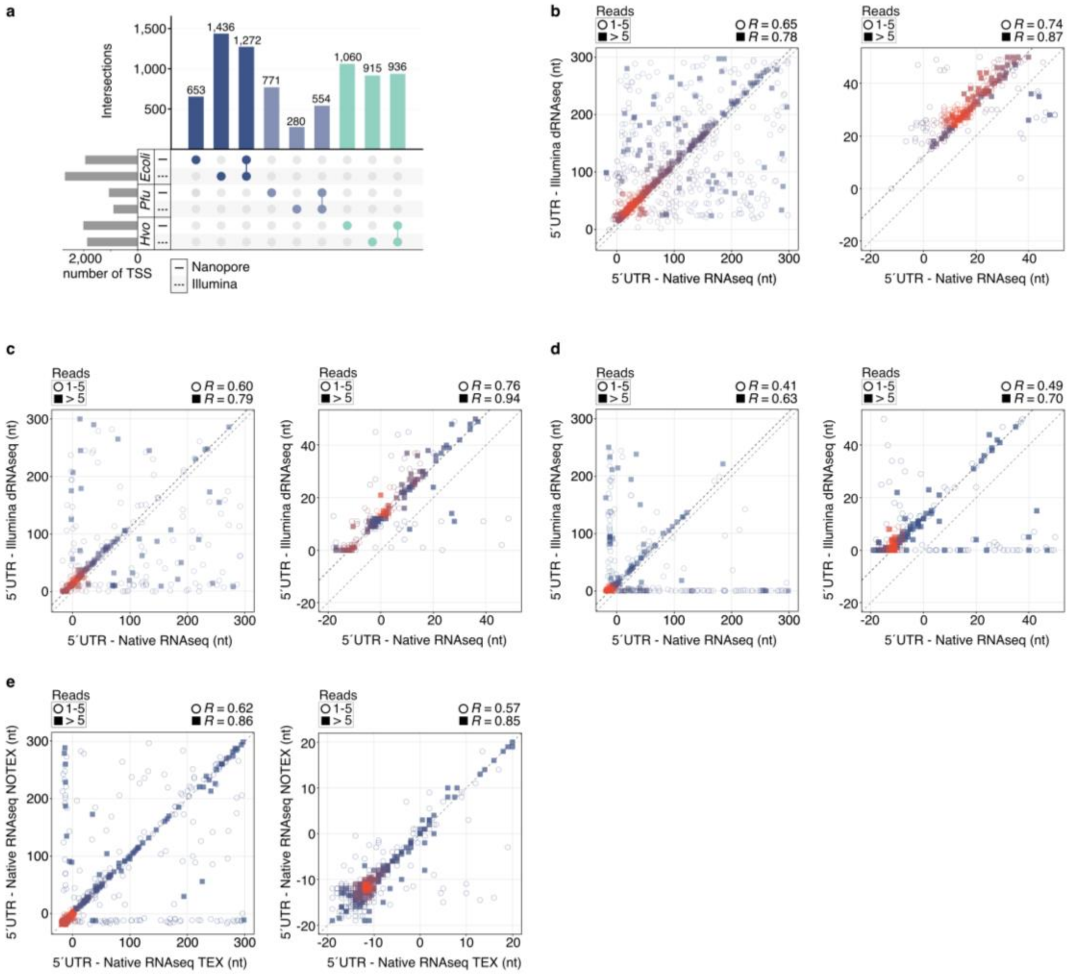


**Supplementary Figure 6 | Transcription start site (TSS) analysis. a**, Primary transcription start sites (TSS) were predicted based on Nanopore reads and compared to Illumina d(ifferential) RNA-Seq data from published data sets for *E. coli*[7], *P. furiosus*[5] and *H. volcanii*[8]. The total number of all genes with a detected TSS is shown as grey barplots and results from the sum of Nanopore-only predicted TSS and the intersection to the Illumina data. Panel **b**-**e**: Correlation analysis between different 5´UTR lenghts with every dot representing one gene, the number of mapped reads indicated by an empty circle (1-5) or filled square (> 5) and density of data shown by color scale from darkblue (low) to high (red). Data are shown for *E. coli* in **b**, *P. furiosus* in **c**, *H. volcanii* in **d**, and TEX and NOTEX set of *H. volcanii* in **e**. Spearman´s rho correlation is shown for all comparisons.

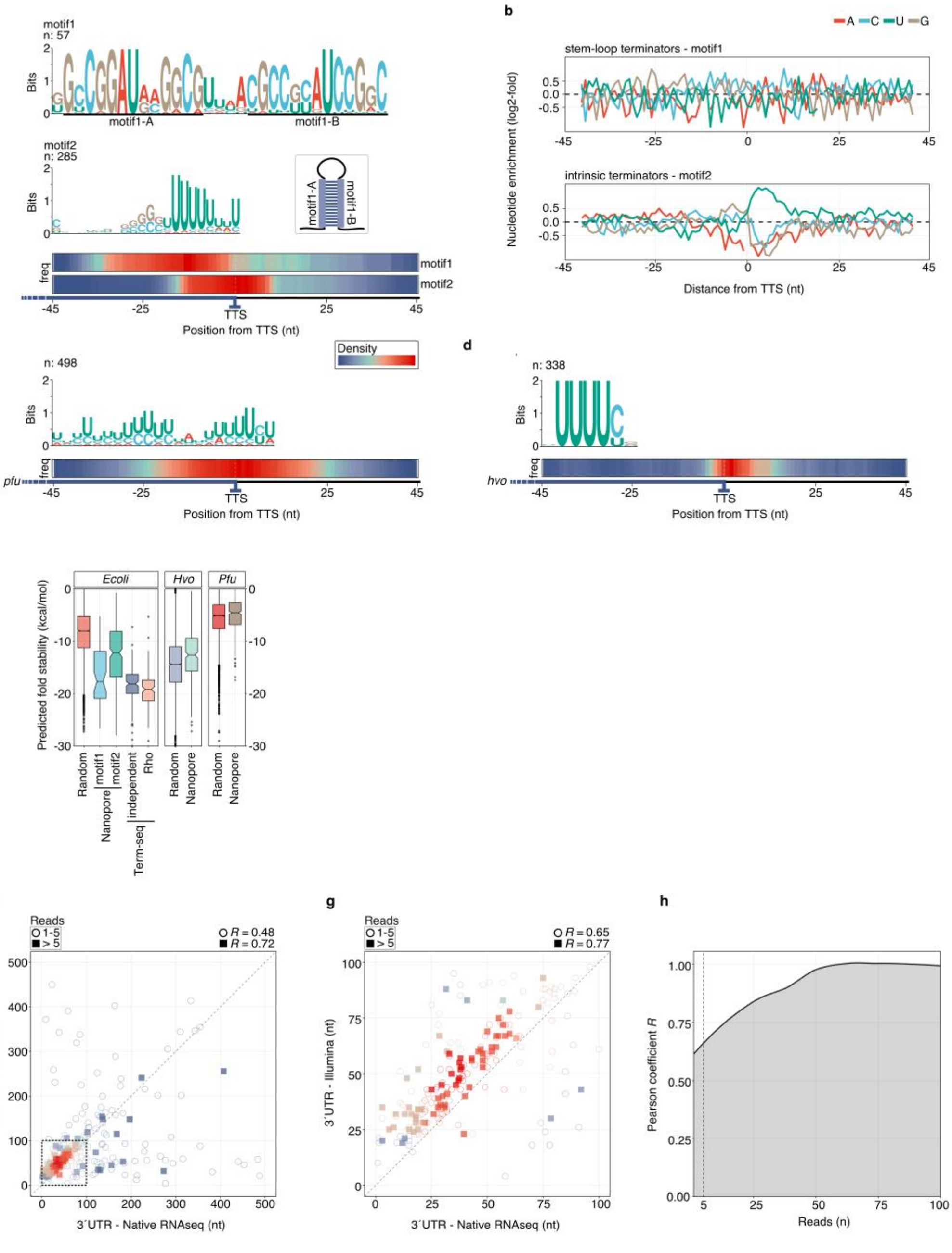


**Supplementary Figure 7 | Transcription termination site (TTS) analysis.** **a**, MEME analysis[9] of TTS in *E. coli* identifies significantly enriched motifs in *E. coli* (stem-loop forming motif1 represent REP sites, intrinsic poly(U)-containing motif2)[10]. Enriched motifs and position of motifs in the scanned region from -45 to +45 from the TTS is shown as sequence pattern and heatmap. **b**, Nucleotide enrichment analysis of terminating regions for motif1 and motif2. Enrichment was calculated by comparing the genomic sequences surrounding the TTS (-45 to +45) to randomly selected intergenic positions (n: 10000). **c**, MEME output (enriched motifs and position-heatmap) of termination site motif scanning in *P. furiosus* and **d**, *H. volcanii*. **e**, Predicted fold stability of selected termination sequences (45 bases upstream of TTS) and comparison to random generated sequences. Fold stability was calculated by RNAfold[11]. **f**, Comparison of 3´UTRs in *H. volcanii* predicted by native RNA sequencing (x-axis) and a short-read Term-Seq approach[12]. Note that every dot represents one gene, the number of mapped reads is indicated by an empty circle (1-5) or filled square (> 5) and density of data is shown by color scale from darkblue (low) to high (red). **g**, Zoom to the majority of 3´UTRs that have 3´UTRs < 100 nts. Spearman´s rho correlation is shown for all comparisons. **h**, Calculated Spearman´s rho coefficient was calculated depending on the number of reads that mapped to a CDS.

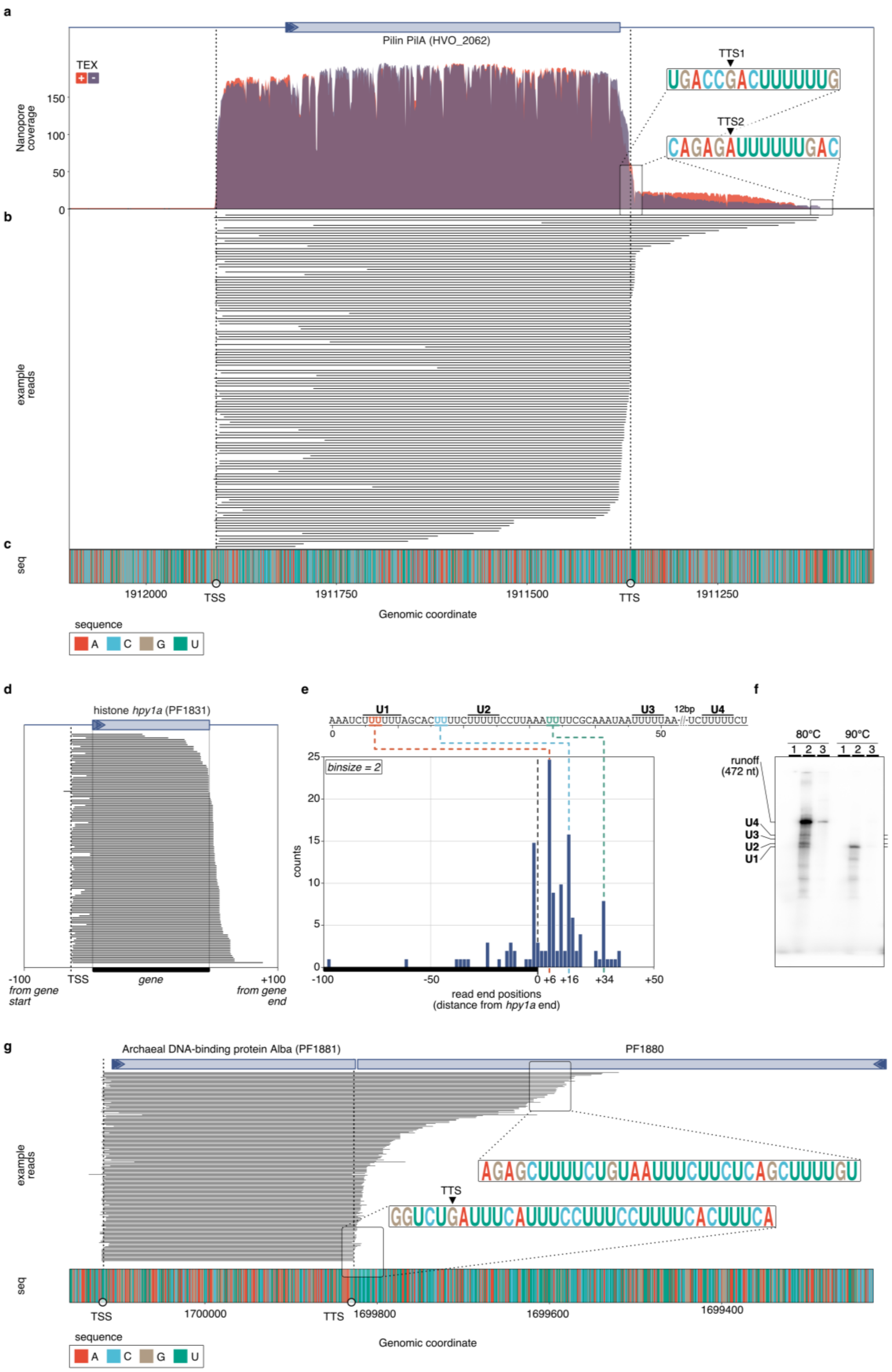


Supplementary Figure 8 | Analysis of termination events for the Pilin PilA gene (HVO_2062) in *Haloferax volcanii* and the hsitone *hpy1a* (PF1831) gene in *P. furiosus* . **a**, Nanopore coverage calculated for both TEX (red) and NOTEX (purple) sample in *Haloferax volcanii.* Coverage drops results from mismatches in the mapped reads. The TU-based estimated transcript boundaries (TSS, TTS) are indicated by vertical dashed lines. Termination specific poly(U) sequences are highlighted in the 3´ UTR region. **b**, IGV snapshot of example reads highlighting different 3´ UTR variants. **c**, Sequence of the plotted region visualized by color-coding the different nucleotides (red: A, blue: C, brown: G, green: U). **d**, Single read track of reads mapping to the *hpy1a* gene. **e**, The 3' read end positions of reads in d were extracted and are shown in a histogram view. Note that one bar reflects the added up counts of 2 positions. In the upper panel the sequence starting from the annotated gene end is shown, with $U_5$ sequences highlighted with U1-U4[13]. Enriched Nanopore read ends are color-coded. **f**, *In vitro* transcription of the *hpy1a* template displayed at 80°C and 90°C using 2mM ATP, 2mM GTP and (1) no CTP, 0.04 mM UTP, (2) 2 mM CTP, 0.04 mM UTP, (3) 2 mM CTP, 2 mM UTP (see Material & Methods). Lenghts of transcripts are depicted on the left. **g**, IGV snapshot of Alba-mapping reads in *P. furiosus*. Two consecutive terminating U-stretches and the color-coded sequence are highlighted.

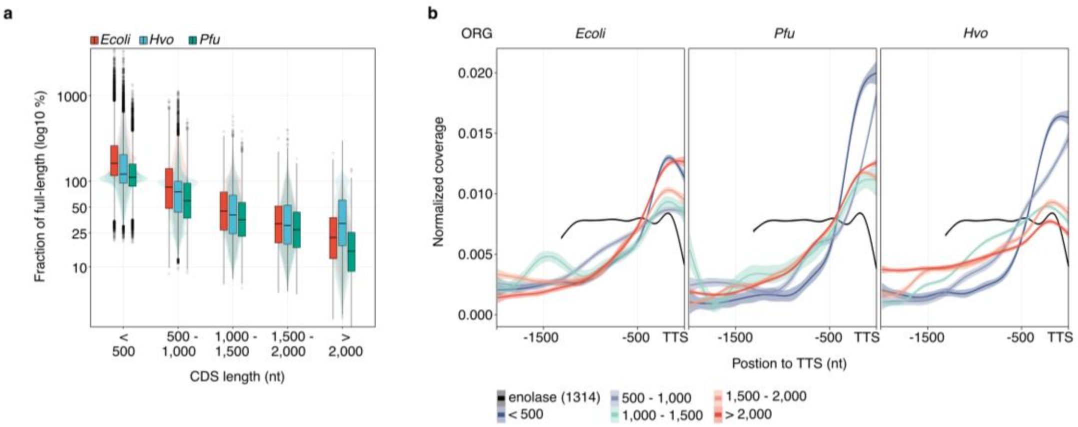


**Supplementary Figure 9 | Prerequisites for transcriptional unit (TU) detection. a**, The fraction of full-length transcripts is compared between protein-coding gene (CDS) classes, that are sorted by gene length in all three TEX-treated samples. **b**, Coverage metaplots aligned to the transcription termination site (TTS) of the 100 most transcribed genes in all samples, show a 3´ to 5´ drop in all native RNA classes (colored by gene length), except for the enolase control (black).

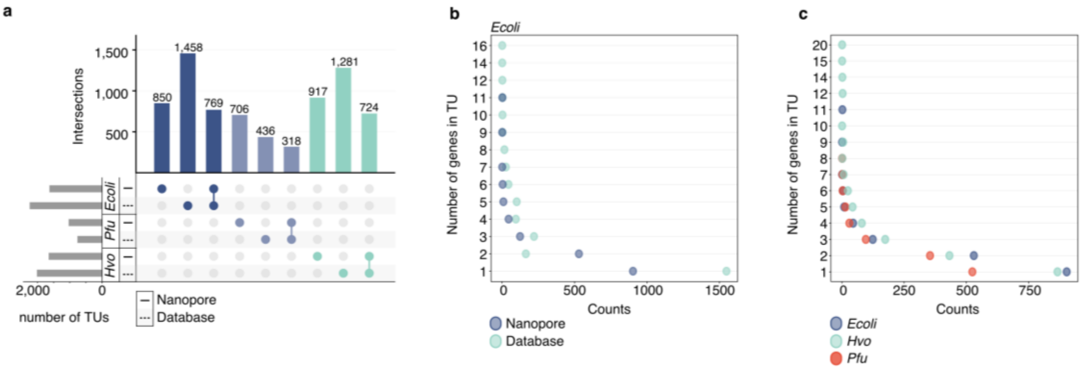


**Supplementary Figure 10 | Detection of transcriptional units (TU) in three prokaryotic model organisms. a**, The number and composition of TUs is compared between Nanopore-detected TUs and operon annotations retrieved from databases for *E. coli*[14], *P. furiosus* (DOOR2 database) and *H. volcanii* (DOOR2)[15]. The total number of TUs is indicated by grey bars and results from the sum of Nanopore-only predicted TUs and the intersection to the database data. **b**, Comparison of the number of genes in transcriptional units in *E. coli* Nanopore data (purple) and the database set (light-green)[14]. **c**, Number of genes in a TU in all three datasets. TU prediction is shown only for TEX-treated samples.

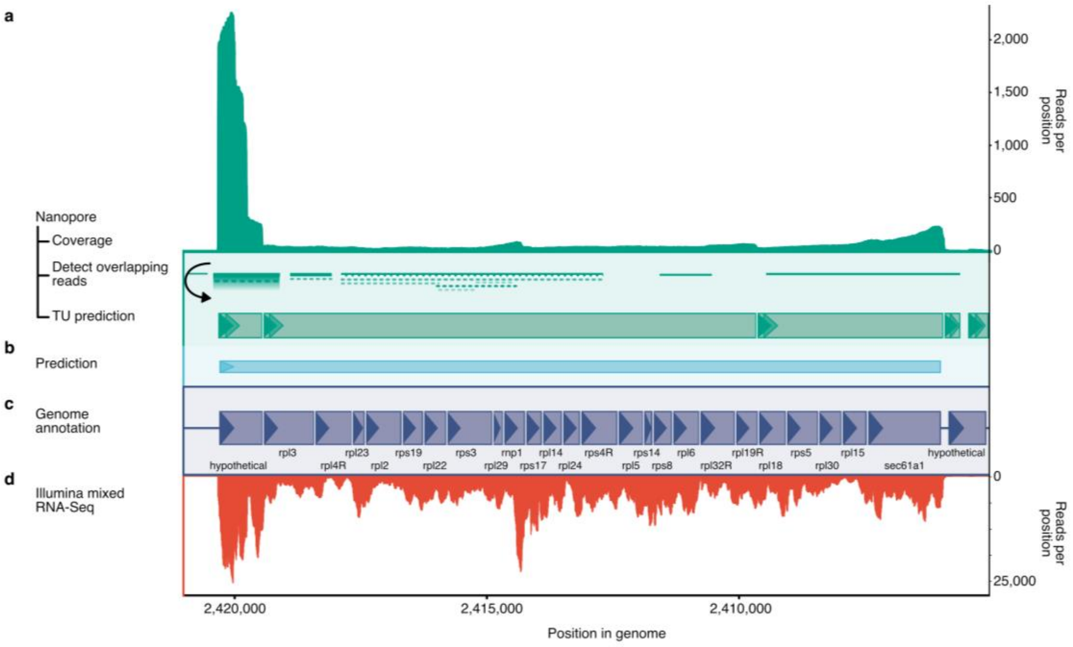


**Supplementary Figure 11 | Large transcriptional unit (TU) annotation of a large ribosomal-protein-containing operon in *Haloferax volcanii*.** **a**, Coverage of Nanopore reads is shown in the top panel. TU prediction is performed by detection and linkage of overlapping reads and splitting them according to a 3´ drop in coverage (compare Supplementary Fig. 8b). Predicted TUs are drawn with green boxes according to scale. **b**, Comparison to bioinformatical prediction using the DOOR2 database[15]. **c**, Genome annotation with abbreviated gene names, boxes drawn to scale and strand indicated by triangles[16]. **d**, Coverage detected using Illumina sequencing of a mixed RNA sample[6].

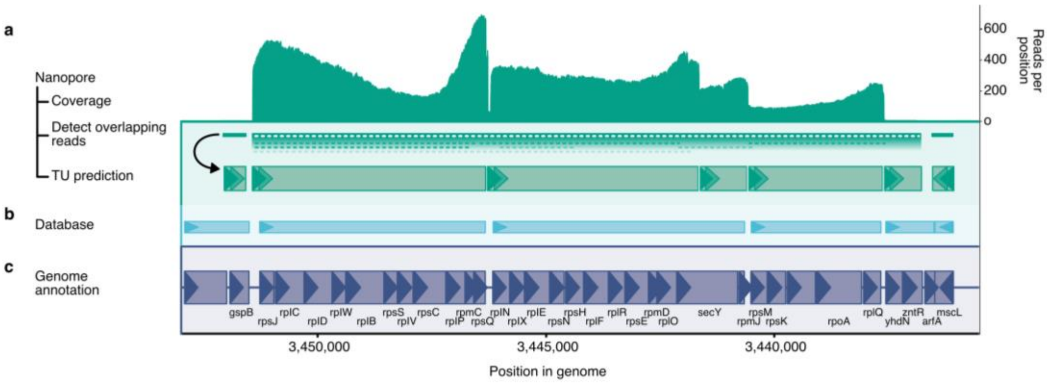


**Supplementary Figure 12 | Transcriptional unit (TU) annotation of the large ribosomal-protein-containing operon in *Escherichia coli*.** **a**, Coverage of Nanopore reads is shown in the top panel. TU prediction is performed by detection and linkage of overlapping reads and splitting them according to a 3´ drop in coverage (compare Supplementary Fig. 8b). **b**, Comparison to published analysis of TUs in *E. coli* based on a bioinformatical approach[14]. **c**, Genome annotation with abbreviated gene names, boxes drawn to scale and strand indicated by triangles[17].

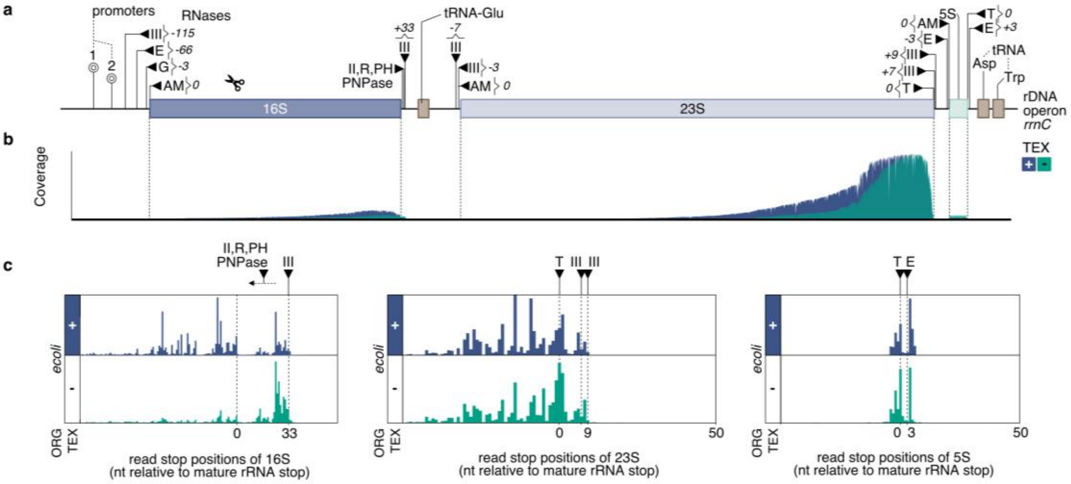


**Supplementary Figure 13 | Detection of ribosomal RNA processing sites in *E. coli*. a**, Transcription of the rDNA locus (rrnC) is starting from two promoters (transcription start sites at -293 and -175)[18]. Precursor RNAs are cleaved by RNases (black triangles) at depicted positions[19–22]. **b**, Histograms of read end positions for 16S, 23S and 5S rRNA. Positions are relative to the annotated boundaries of mature rRNAs and shown for TEX (+, purple) and NOTEX (-, green) samples.

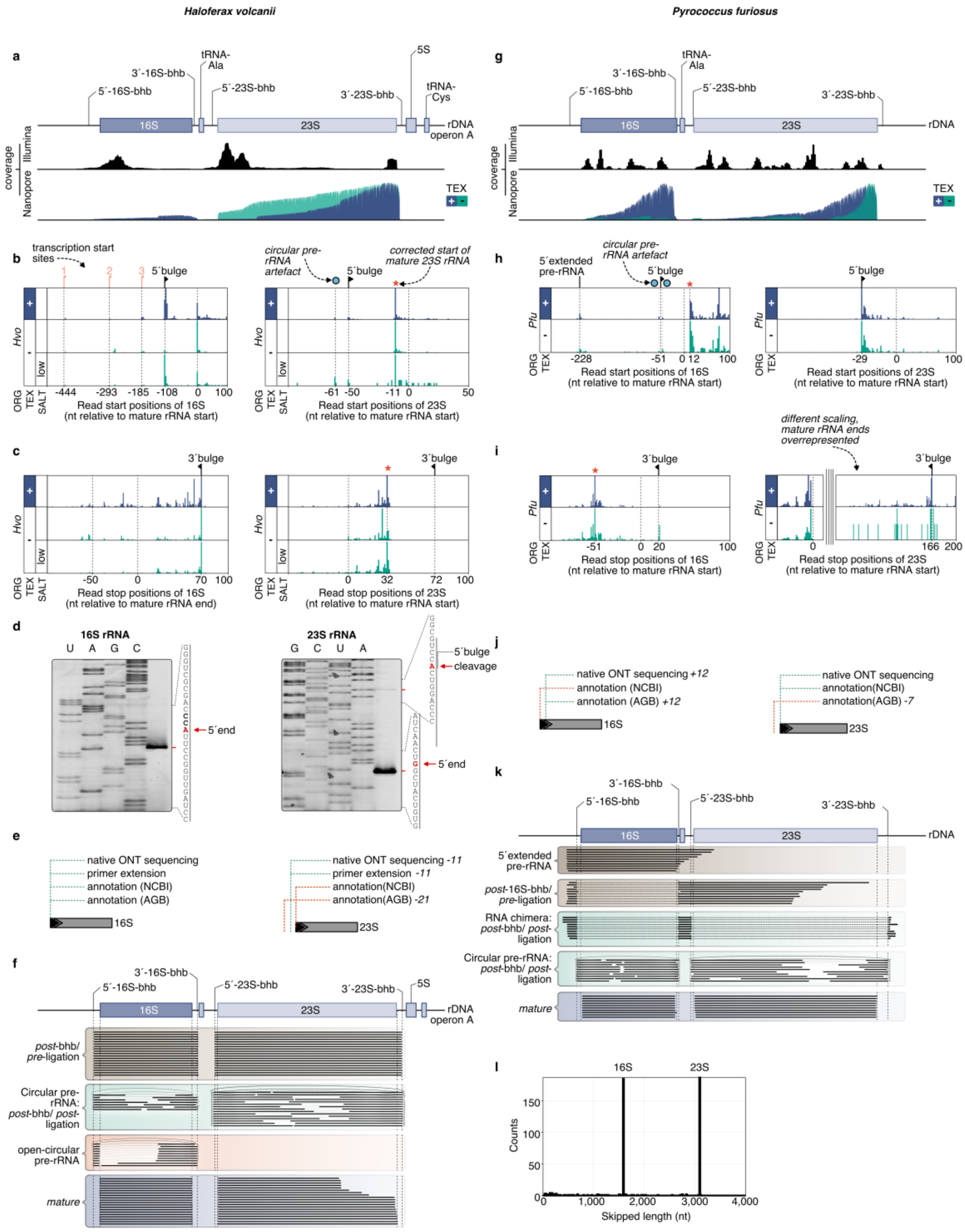


**Supplementary Figure 14 | rRNA processing site detection in *H. volcanii* and *P. furiosus*. a/g**, Schematic of the rDNA locus operon A in *H. volcanii* and the only rDNA locus in *P. furiosus*. Nanopore coverage tracks are shown for TEX (+, blue) and NOTEX (-, green) samples and compared to short-read Illumina coverages. **b/h**, Histograms of read start positions and **c/i**, read end positions of 16S and 23S rRNA relative to annotated boundaries of mature rRNAs of TEX (+) and NOTEX (-) samples. Cleavage sites are indicated by black triangles. Asterisks mark sites of mature rRNA that are potentially not annotated correctly. **d**, Mapping of mature 16S and 23S rRNA 5´ends by primer extension. Primer extension were performed with the indicated primers as described in Material and Methods. From the comparison to a sequencing ladder, positions of mature 16S rRNA and 23S rRNA and the 5´bulge of the 23S rRNA could be assigned (red arrows). **e/j** Annotation of the 16S and 23S rRNA start based on NCBI (https://www.ncbi.nlm.nih.gov/genome/) and archaeal genome browser (AGB, http://archaea.ucsc.edu) versions, compared to positions enriched in native ONT sequencing and experimentally verified by primer extension. **f/k** Single-read tracks of read categories derived from (i) co-occurrence analysis of read start and stop positions, (ii) the number of junctions and (iii) clipping properties. **l**, Skipped length of the RNA chimera class verifies accuracy of “spliced” read detection.

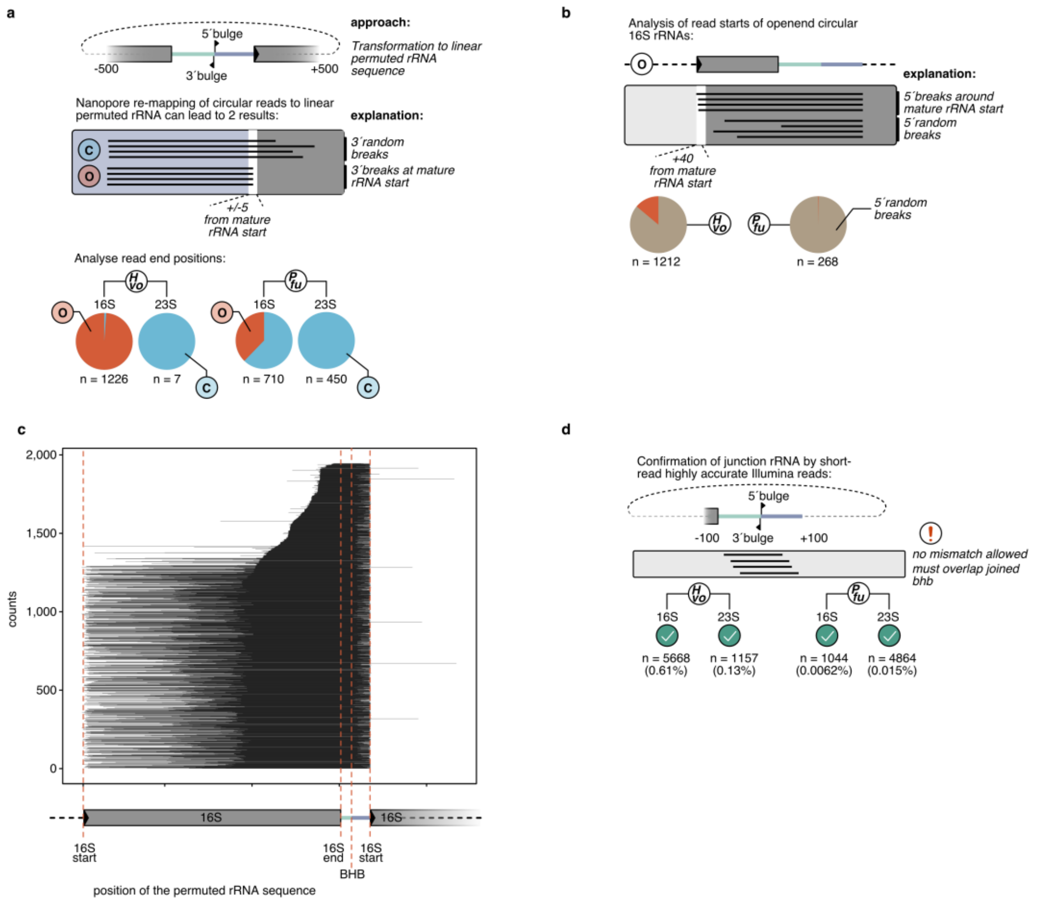


**Supplementary Figure 15| Circular read detection of archaeal rRNA precursors. a**, Circular reads were confirmed by re-mapping Nanopore reads to a permuted rRNA sequence, containing the joined bulge-helix-bulge site. We categorized reads based on their 3´terminal positions as circular pre-RNA (C, random breaks) and opened-circular pre-rRNAs (O, 3´end at mature rRNA start) and counted the number of reads fulfilling the criteria (*Pfu* TEX and *Hvo* NOTEX). **b**, Read start positions of opened-circular pre-rRNAs. **c**, Single read plotting of *H. volcanii* (NOTEX) sample reads mapping to the permuted 16S rRNA sequence. Note that the 5´ends have been extended by their left clipping size, so the full 16S gene is covered. **d**, Circular reads were confirmed by mapping highly accurate Illumina reads to the permuted sequence, allowing for no mismatch and filtering out all reads that do not overlap the joined bulge-helix-bulge.

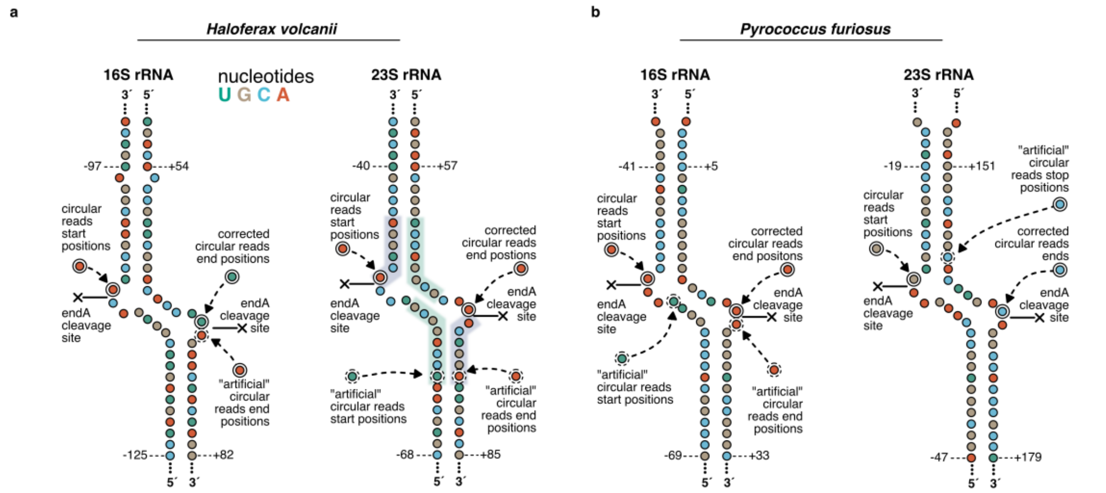


**Supplementary Figure 16 | Secondary structure prediction of bulge-helix-bulges. a**, Secondary structure predicted was performed by RNAfold from the Vienna RNA package[11]. Artificial circular read end and start positions are caused by similarities in the 5´leading and 3´-trailing sequences, but are mostly accurately detected at the endA cleavage site in *H. volcanii* and **b**, *P. furiosus*.

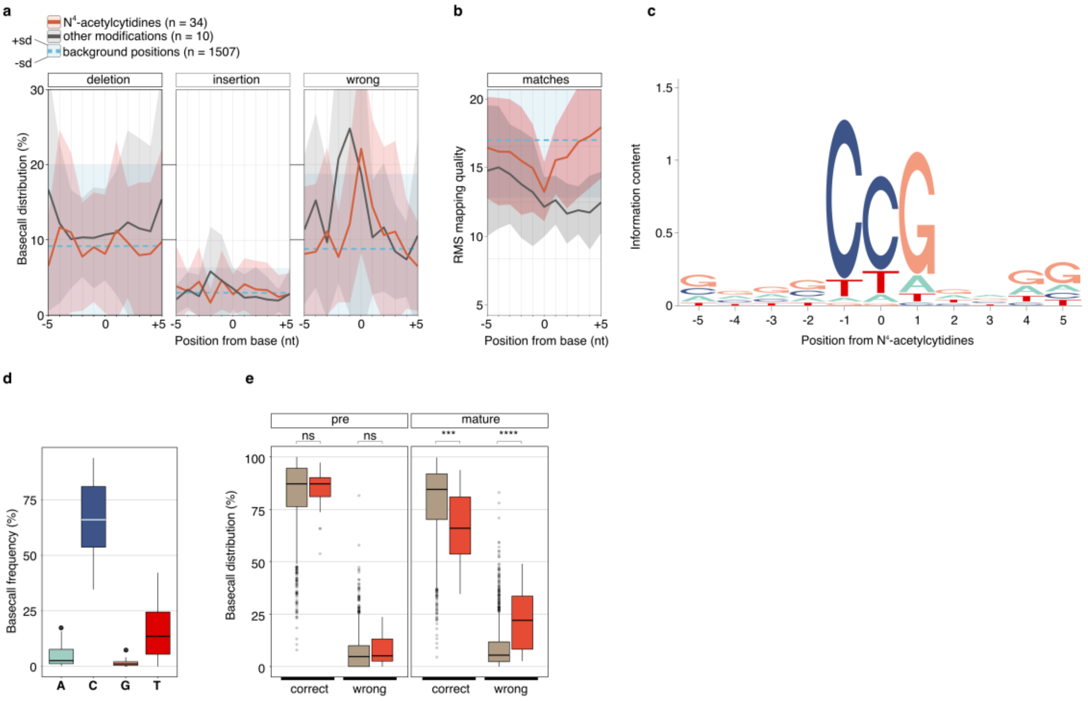


**Supplementary Figure 17 | Detection of N⁴-acetylcytidine modifications in *P. furiosus*.** **a**, The proportion of deletions, insertions and wrong bases is shown in a window from -5 to +5 from the presumably modified or background base. Shaded areas show the upper and lower standard deviation, while the lines show the mean values of N⁴-acetylated positions (red), diverse other modifications (grey) and all other positions of the 16S rRNA in *P. furiosus* using the recently established 16S rRNA modification pattern in the close relative *Pyrococcus abyssi*[23]. **b**, Root mean square (RMS) mapping quality in the surrounding sequence context of potentially modified bases. **c**, Sequence logo of actual sequenced bases averaged for all 34 potentially N⁴-acetylated positions in the 16S rRNA. **d**, Frequency of each nucleotide to be basecalled in a N⁴-acetylated CCG context. **e**, Comparison of the basecall properties of 5´-extended pre-rRNA and mature rRNAs. Statistical significance (p-values, T-test) is indicated by asterisks (p-value > 0.05: ns (not significant), p <= 0.0001: ****).

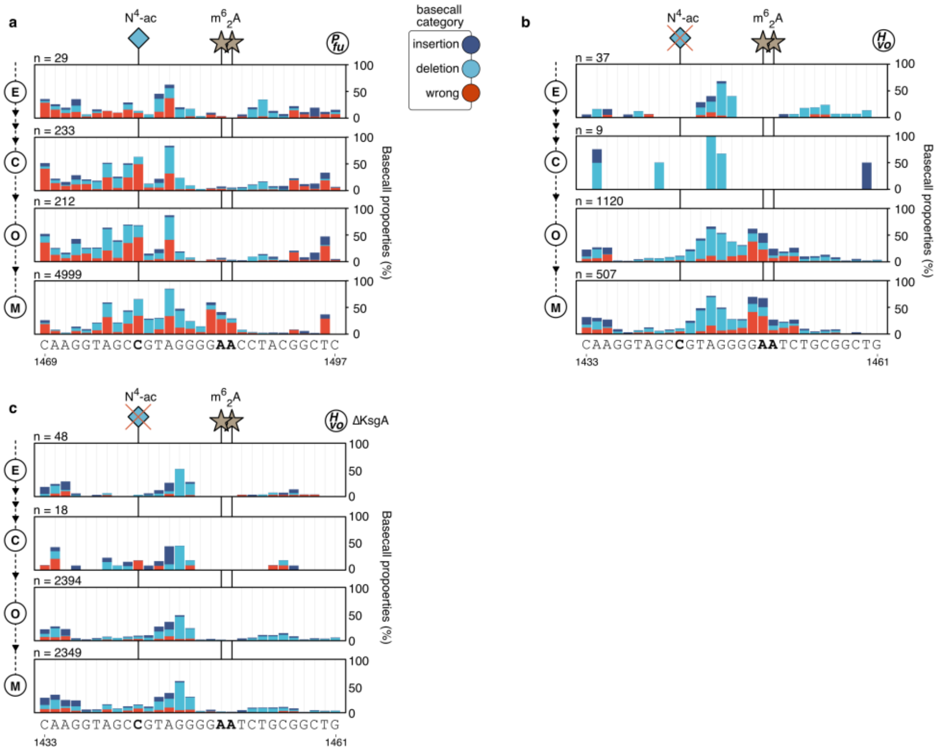


**Supplementary Figure 18 | Detection of h45-N⁴-acetylation and KsgA-dependent $m^6_2A$ modification. a**, Basecall properties (insertion: dark-blue, deletion: light-blue, wrong: red) are shown for read subsets reflecting different stages of rRNA maturation (E: 5´-extended pre-rRNAs, C: circular pre-rRNAs, O: opened-circular pre-rRNAs, M: mature rRNAs) in *P. furiosus*, **b**, *H. volcanii* wildtype and **c**, *H. volcanii* ΔKsgA.
